# A deep phenotyping study in mouse and iPSC models to understand the role of oligodendroglia in optic neuropathy in Wolfram syndrome

**DOI:** 10.1101/2024.04.28.591501

**Authors:** K. Ahuja, M. Vandenabeele, F. Nami, E. Lefevere, J. Van hoecke, S. Bergmans, M. Claes, T. Vervliet, K. Neyrincki, T. Burg, D. De Herdt, P. Bhaskar, Y. Zhu, Z. J. Looser, J. Loncke, W. Gsell, M. Plaas, P. Agostinis, J.V. Swinnen, L. Van Den Bosch, G. Bultynck, A. S. Saab, E. Wolfs, Y.C. Chai, U. Himmelreich, C. Verfaillie, L. Moons, L. De Groef

## Abstract

Wolfram syndrome (WS) is a rare childhood disease characterized by diabetes mellitus, diabetes insipidus, blindness, deafness, neurodegeneration and eventually early death, due to autosomal recessive mutations in the *WFS1* (and *WFS2*) gene. While it is categorized as a neurodegenerative disease, it is increasingly becoming clear that other cell types besides neurons may be affected and contribute to the pathogenesis. MRI studies in patients and phenotyping studies in WS rodent models indicate white matter/myelin loss, implicating a role for oligodendroglia in WS-associated neurodegeneration. In this study, we sought to determine if oligodendroglia are affected in WS and whether their dysfunction may be the primary cause of the observed optic neuropathy and brain neurodegeneration. We demonstrate that 7.5-month-old *Wfs1*^Δexon8^ mice display signs of abnormal myelination and a reduced number of oligodendrocyte precursor cells (OPCs) as well as abnormal axonal conduction in the optic nerve. An MRI study of the brain furthermore revealed grey and white matter loss in the cerebellum, brainstem, and superior colliculus, as is seen in WS patients. To further dissect the role of oligodendroglia in WS, we performed a transcriptomics study of WS patient iPSC-derived OPCs and pre-myelinating oligodendrocytes. Transcriptional changes compared to isogenic control cells were found for genes with a role in ER function. However, a deep phenotyping study of these WS patient iPSC-derived oligodendroglia unveiled normal differentiation, mitochondria-associated endoplasmic reticulum (ER) membrane interactions and mitochondrial function, and no overt signs of ER stress. Overall, the current study indicates that oligodendroglia functions are largely preserved in the WS mouse and patient iPSC-derived models used in this study. These findings do not support a major defect in oligodendroglia function as the primary cause of WS, and warrant further investigation of neurons and neuron-oligodendroglia interactions as a target for future neuroprotective or -restorative treatments for WS.

## 1. Introduction

Wolfram syndrome (WS) type 1 is a rare hereditary disease affecting 1 in 770.000 newborns that, in most cases, is caused by recessive mutations in the Wolfram syndrome 1 (*WFS1)* gene [23, 69]. The disease symptoms are often referred to as ‘DIDMOAD’, an acronym of diabetes insipidus, diabetes mellitus, optic atrophy and deafness. However, patients may also experience a range of other (neurological) symptoms, including ataxia, urinary tract problems, psychiatric problems, apnea and autonomic dysfunction [12, 29]. The disease manifests in early childhood and gradually worsens until patients die around the age of 30 years [69]. Given the monogenic cause of the disease, research efforts have focused on elucidating the role of the WFS1 protein in WS pathogenesis. In Wolfram patients, loss-of-function and missense mutations lead to a complete absence or a decreased stability of the WFS1 protein, suggesting that WS is caused by a loss of WFS1 function [19, 23]. WFS1 encodes for an endoplasmic reticulum (ER) membrane protein that resides at the mitochondria-associated ER membranes (MAMs) [12, 35]. There, it participates in multiple signalling cascades: WFS1 regulates the ER stress response [15], steady-state ER Ca^2+^ concentrations [67, 74], Ca^2+^transfer between the ER and mitochondria [3], and mitochondrial dynamics [8, 9, 24, 75]. It is proposed that WFS1 deficiency partakes in the unfolded protein response pathway triggered by ER stress, which, if unresolved, initiates apoptotic pathways leading to cell demise.

Notably, until today, research has mainly focused on the diabetic phenotype of WS and the aforementioned WFS1 functions have mostly been studied in the islets of Langerhans and β-cells of the pancreas [17, 26]. However, diabetes has become a chronic illness that can be managed via drug therapy and only has a limited impact on the quality of life of Wolfram patients. This contrasts with the neurodegenerative phenotype of the disease, which is devastating and cannot be controlled by any means. In recent years, efforts to improve our knowledge on the neuronal phenotype of Wolfram patients and *Wfs1* animal models ensued. In neurons, WFS1 deficiency has been associated with MAM dysregulation, impaired neurite outgrowth and synapse formation, and increased cytosolic Ca^2+^ concentration [54, 75]. A recent study using brain organoids showed that this abnormal synapse formation and function goes hand in hand with decreased expression of the excitatory amino acid transporter 2 (EAAT2) and compromised glutamate clearance in astrocytes [73]. MRI studies in Wolfram patients found white matter changes and volume loss in different brain regions, most prominently in the brainstem, cerebellum and optic radiations, already early in the disease pathogenesis [18, 41]. In addition, optic nerve and brainstem degeneration, as well as optic nerve demyelination, have also been observed in *Wfs1* rat and mouse models [9, 53]. In the *Wfs1*^Λ1exon8^ mouse model, disturbances in the dopaminergic and GABAergic system, together with changes in motor function and anxiety have been described, thereby mirroring the motor and psychiatric problems seen in patients [42, 56, 71]. Furthermore, a recent study by Rossi *et al.* reported that retinal degeneration and vision loss in the *Wfs1*^Λ1exon8^ mouse model is preceded by loss of glial homeostasis, MCT1-dependent metabolic dysfunction and myelin derangement in the optic nerve [59]. While these studies all find a clear neurodegenerative phenotype, they also show white matter defects, hence raising the question whether the neurodegenerative component of WS could be mainly driven by an oligodendroglia rather than a neuronal pathology. Oligodendrocytes, due to their continuous, high production of myelin, are particularly sensitive to ER stress [33], and their dysfunction or loss will have detrimental consequences for the highly energy-demanding neurons. In that case, neurodegeneration in WS might relate to the loss of the supportive functions that are being fulfilled by oligodendroglia in the central nervous system (CNS). Indeed, oligodendrocytes physically protect and increase conductivity by insulating axons with myelin sheaths, but oligodendroglia also provide metabolic support to axons by transferring metabolites, like lactate or pyruvate, to fuel the axonal energy production [48, 60], and they support the formation and function of neuronal synapses and neural circuits [46, 68]. Thus, dysfunction or loss of oligodendroglia might substantially contribute to neurodegeneration.

In this study, we combined a neuropathological analysis of the retina, optic nerve and brain of a WS mouse model with a deep phenotyping study of WS patient iPSC-derived oligodendroglia, to disentangle the role of the oligodendroglia in WS pathogenesis. Our investigations reveal that the *Wfs1* KO mouse presents with grey and white matter abnormalities in the brain, and abnormal axonal conduction combined with subtle thinning of the myelin sheath and a reduction of the oligodendrocyte precursor (OPC) pool in the optic nerve. Next, a differential transcriptomics study of oligodendroglia (i.e., OPCs and premyelinating oligodendrocytes) derived from WS patient versus isogenic control iPSCs led to the identification of several genes that have previously been linked with WS-like syndromes characterized by impaired vision and hearing, as well as cellular processes previously linked to WFS1 deficiency, such as ER function. However, *in vitro* assays using WS patient iPSC-derived oligodendroglia did not uncover any signs of ER stress, abnormal ER-mitochondria interactions, or mitochondrial dysfunction. Hence, we conclude that cell-intrinsic defects in oligodendroglia are minor and unlikely to be the primary cause of WS, and that future research should focus on the neurons and their intricate interactions with oligodendroglia that govern CNS function and integrity.

## 2. Material and Methods

### 2.1. Experimental animals

All experiments were performed using *Wfs1*^Δexon8^ mice (*Wfs1* KO mice) and wild type littermates (129S6/SvEvTac and C57BL/6 mixed background) of both sexes, at the age of 3, 4.5, 6 or 7.5 months. This mouse model was originally created by Luuk *et al.* and has a disrupted exon 8 in the *Wfs1* gene [42]. Mice were bred under standard laboratory conditions.

#### Optical coherence tomography

Upon general anesthesia and pupil dilatation (0.5% tropicamide), optical coherence tomography scans of the retina (1000 A-scans, 100 B-scans, 1.4 x 1.4 mm) (Envisu R2200, Bioptigen) were acquired. Retinal layer thickness was measured using InVivoVue Diver 3.0.8 software (Bioptigen), at 16 locations equally spaced around the optic nerve head and averaged per mouse, all as described in [70].

### 2.2. Optomotor response test

Visual performance was tested using the virtual reality optomotor test (Optomotry, Cerebral Mechanics), as previously described [65, 70]. Briefly, the mouse was placed on a platform in the testing arena and vertical black-white sine-wave gratings were projected on the screens. Spatial frequency thresholds were measured for each eye under 100% contrast, via a simple staircase procedure. The highest spatial frequency that the mouse could track was identified as the visual acuity. For contrast sensitivity, a similar approach was used, but contrast was systematically reduced until the contrast threshold was identified. Contrast threshold was identified at three spatial frequencies (0.103, 0.192, 0.272 cyc/deg), and calculated as described in [55].

### 2.3. Electroretinograms and visual evoked potentials

Following overnight dark adaptation, electroretinograms (Celeris, Diagnosys) were recorded for anesthetized mice as described in [11]. Lens electrodes with integrated stimulators were placed on the cornea after pupil dilation (0.5% tropicamide and 15% phenylephrine), and a needle scalp electrode was inserted above the midline near the visual cortex. The ground and reference electrode were placed in the tail base and cheek, respectively. Eyes were alternately stimulated, and full field visual evoked potential responses were recorded at a single flash intensity of 0.05 cd*s/m^2^, averaging 300 brief flashes with an inter-sweep delay of 690 ms. To measure the scotopic threshold response, the responses from 50 light flashes with a single-flash intensity of 0.0001 cd·s/m^2^ and an inter-sweep delay of 1 s were averaged. Data were analyzed with Espion v6.65.1 software (Diagnosys) according to [11]. Visual evoked potential and scotopic threshold response amplitudes were defined as the amplitude from the baseline to the trough of the negative visual evoked potential response, and from the baseline to the peak of the positive scotopic threshold response, respectively.

### 2.4. Ex vivo compound action potentials

Acute optic nerve preparations for compound action potential (CAP) recordings were carried out as previously described [36]. In brief, optic nerves were isolated after isoflurane anesthesia and decapitation and placed in an interface perfusion chamber (Haas Top, Harvard Apparatus), perfused with artificial cerebrospinal fluid (ACSF containing in mM: 126 NaCl, 3 KCl, 2 CaCl_2_, 1.25 NaH_2_PO_4_, 26 NaHCO_3_, 2 MgSO_4_, 10 glucose, pH 7.4) at 37°C using a TC-10 temperature control system (npi electronic), and continuously oxygenated with 95% O_2_ and 5% CO_2_. Nerve ends were inserted into custom-made suction electrodes filled with ACSF and stimulated evoking the CAP. The optic nerve was first stimulated at 0.4 Hz for 1 min to obtain baseline values, followed by a stepwise increase in stimulation frequency, with intervals of 1 min at frequencies of 1, 10, 25, and 50 Hz and a recovery period of 10 min with 0.4 Hz.

### 2.5. Magnetic resonance imaging

Mice were anaesthetized with isofluorane (2.5% induction, 1-1.5% for maintenance). Body temperature was kept constant at 37 ± 1 °C, respiration rate was monitored during scanning and kept above 80 min^-1^ by adjusting isoflurane concentrations. MR images were acquired using a 9.4 Tesla preclinical MR scanner with a horizontal bore of 20 cm (Biospec 94/20, Bruker Biospin) and equipped with an actively shielded gradient set of up to 600 mT m^-1^. Brain images were acquired using a linearly polarized resonator (transmit) in combination with an actively decoupled surface coil for the mouse brain (Bruker Biospin) as previously described [64]. For the acquisition of manganese enhance MRI, mice received a unilateral intravitreal injection of 2 μl of 0.5 M MnCl_2_ 20 to 24 hours before image acquisition, similar to previous reports [34].

MRI was performed at the age of 3, 4.5, 6 and 7.5 months. After initial localizer images, T_2_-weighted two-dimensional (2D) MRI scans of the brain were acquired in axial orientation using a spin-echo sequence (rapid acquisition with relaxation enhancement, RARE). Acquisition parameters: repetition time (TR) = 2500ms, echo time (TE) = 33ms, RARE factor (RF) = 8, 20 slices, 0.7mm slice thickness, 70 µm in plane resolution and number of averages (NA) = 1. For the analysis of Mn^2+^ distribution T_1_-weighted MRI were acquired using a 3D gradient echo sequence (FLASH) with TR=30 ms, TE=5 ms, 60° pulse, NA = 2, field of view (FOV)= 2.0 x 1.5 x 1.5 cm and an isotropic spatial resolution of 156 µm. T_1_ maps were determined using a spin echo multiple TR sequence (RAREVTR) with TE = 20 ms, RF = 8, NA = 2, TR = 500, 750, 1500, 2500, 6000 ms and geometric parameters as for the 2D with an in-plane resolution of 125 µm. Finally, diffusion MRI was acquired for the calculation of apparent diffusion coefficients (ADC), mean diffusivity (MD) and fractional anisotropy (FA). Parameters for diffusion MRI were as following: TR = 750 ms, TE = 20 ms, NA = 1, spin echo readout, 6 directions, b-values = 0, 100, 600, 1200 s/mm^3^, and geometric parameters as for the 2D with an in-plane resolution of 125 µm. The Bruker software ParaVision was used for image acquisition, processing and calculation of parametric maps. Examples for the respective images are shown in supplementary figure S1.

### 2.6. Human induced pluripotent stem cell culture

Two WS patient derived induced pluripotent stem cell (iPSC) lines (kindly provided by Prof. F. Urano, Washington University School of Medicine), and their genetically corrected isogenic controls made by adenosine base editing, have been used in this study: patient 1, corresponding to homozygous point mutation at c.2002, and patient 2, referring to compound heterozygous mutation at c.376 and c.1838 [47]. Both patient and isogenic iPSC lines were cultured on human Matrigel (VWR)-coated 6-well plates (Corning) in E8 basal medium (Gibco) complemented with E8 supplement Flex (Gibco) and 5 U/ml Penicillin-Streptomycin (Gibco), and passaged two times a week using 0.5 mM EDTA (Gibco) (in PBS). Routine checkups for mycoplasma contamination were done using MycoAlert Mycoplasma Detection Kit (Lonza) to ensure data reliability.

### 2.7. Differentiation of iPSCs to OPCs and pre-myelinating oligodendrocytes

All iPSC lines were differentiated to OPCs and pre-myelinating oligodendrocytes (OPCs/pmOLs), using overexpression of the transcription factor *SOX10*, as described in Garcia-Leon *et al*. [16]. *SOX10* overexpression iPSCs were dissociated with accutase (Sigma) and plated at 25,000 cells/cm^2^ in a 6-well plate. Oligodendrocyte induction was done by adding neurobasal medium supplemented with 0.1 μM RA (Sigma), 10μM SB431542 (Tocris), and 1μM LDN-193189 (Milteny) for the next 6 days. From day 7 on, medium was replaced with neurobasal medium containing 0.1 μM RA, and 1μM SAG (Milllipore) for 3 days. The cells were then dissociated with accutase (Sigma) and replated at 50000-75000 cells/cm^2^ in a poly-L-ornithin-laminin coated 6-well plate with neurobasal medium supplemented with 10 ng/ml PDGFaa (Peprotech), 10 ng/ml IGF1 (Peprotech), 5ng/ml HGF (Peprotech), 10 ng/ml NT3 (Peprotech), 60 ng/ml T3 (Sigma), 100 ng/ml Biotin (Sigma), 1 μM cAMP (Sigma), and 5 μg/ml doxycycline (Sigma) (from here on referred to as oligodendrocyte maturation medium (OMM)) to induce oligodendrocyte fate, until day 23. On day 24, the cells were dissociated with accutase and used for experiments or cryopreserved in liquid nitrogen for future use.

### 2.8. RT-qPCR

Total RNA was isolated and purified using the Quick-RNA Microprep Kit (Zymo research), and cDNA preparation was done with the SuperScript^TM^ III First-Strand Synthesis System kit (Thermofisher), on 500ng-1ug RNA. RT-qPCR for oligodendrocyte lineage markers was performed using Platinum SYBR Green qPCR Supermix-UDG (Thermofisher). The sequence of the primers is listed in supplementary table S1.

### 2.9. X-Box-Binding protein 1 (XBP1) splicing assay

RNA isolation was performed as described above, with the Quick-RNA Microprep Kit (Zymo research) and SuperScript^TM^ III First-Strand Synthesis System kit (Thermofisher). Next, 140 ng cDNA was used to set-up a PCR reaction using Platinum® Taq mix (Invitrogen™) (Supplementary Table 1). The PCR product was run on 2.5% agarose/1x TAE gel with SYBR Safe. In addition to the spliced and unspliced amplicon, a hybrid amplicon was also seen in the gel. Quantification was done as the ratio of spliced amplicon (spliced amplicon + 50% of the hybrid amplicon) and total XBP1 amplicon (spliced + unspliced + hybrid amplicon) using ImageJ [58].

### 2.10. Real time cell metabolic analysis

OPCs/pmOLs were plated on a Seahorse XF24 (Agilent) Extracellular Flux Analyzer culture plate in OMM supplemented with 5 µg/ml doxycycline (Sigma) and RevitaCell supplement (Gibco). The next day, medium change with Mito Stress assay medium (Agilent) containing 10 mM glucose, 2 mM glutamine and 1 mM sodium pyruvate was done one hour before the Mito Stress assay. Addition of 1 μM oligomycin (Sigma-Aldrich), 2.0 μM FCCP (Sanbio) and 0.5 μM antimycin A (Sigma-Aldrich), allowed to detect the mitochondrial respiration activity. After the assay, cells were detached using accutase and counted with an automated cell counter (Nucleocounter 900-002, Chemometec) to enable normalization.

### 2.11. Intracellular Ca^2+^ measurements

OPCs/pmOLs were plated on a 4-chambered glass bottom dish (Cellvis) in OMM supplemented with 5 µg/ml doxycycline (Sigma) and RevitaCell (Gibco). After 2 days, Ca^2+^ recording was performed by loading 1.25 µM Cal-520AM (Abcam, ab171868) in OMM for one hour. PBS washing was performed twice, followed by addition of modified Krebs-Ringer solution (135 mM NaCl, 6.2 mM KCl, 1.2 mM MgCl_2_, 12 mM HEPES, pH 7.3, 11.5 mM glucose and 2 mM CaCl_2_) for imaging. For imaging (eclipse Ti2, Nikon), a baseline was recorded for 30 s followed by addition of different stimulants (10 μM ATP or 10 μM acetylcholine). Cal-520 was excited at 480 nm, after which fluorescent intensity alterations were measured at 510 nm. Area under the curve and peak intensity were quantified using ImageJ software. Fluorescent signals were plotted as F/F_0_, with F_0_ the mean Cal-520AM fluorescent intensity from the first 10 s of the baseline measurement [49].

### 2.12. Proximity ligation assay

The proximity ligation assay was done as per manufacturer’s protocol, using the Duolink^®^ Proximity Ligation Assay kit (Sigma-Aldrich). Briefly, OPCs/pmOLs were plated on a 16 well CultureWell™ Chambered Coverglass (Invitrogen) in OMM supplemented with 5 µg/ml doxycycline (Sigma) and RevitaCell (Gibco). After 2 days, the cells were fixed with 4% paraformaldehyde (PFA) (Thermo Fisher) for 10 min, permeabilized with 0.1% Triton X100 in PBS and incubated overnight with the primary antibodies diluted in blocking serum (anti-protein tyrosine phosphatase interacting protein 51 (PTPIP51) (Abcam, ab224081) and anti-vesicle-associated membrane protein B (VAPB) (RnD Systems, MAB58551)) at 4°C. Secondary antibodies were added to the cells in blocking serum for 1 hour at 37°C. Further ligation and amplification were performed by adding ligation buffer for 90 min, and amplification buffer for 100 min, at 37°C. Lastly, slides were mounted with Duolink *In Situ* Mounting Medium containing 4’,6-diamidino-2-phenylindole (DAPI). Primary antibodies were omitted as a negative control. Quantification of VAPB-PTPIP51 interactions was done on confocal microscopy images (C2, Nikon), using ImageJ software, as described [38].

### 2.13. Lipidomics

Lipids were extracted from a pellet of 500,000 cells, homogenized in 700 μl water with a handheld sonicator, and mixed with 800 μl 1 N HCl:CH_3_OH 1:8 (v/v), 900 μl CHCl_3_ and 200 μg/ml 2,6-di-tert-butyl-4-methylphenol (BHT; Sigma-Aldrich), 3 μl of SPLASH® LIPIDOMIX® Mass Spec Standard (Avanti Polar Lipids, 330707), and 3 μl of Ceramides and 3 μl of Hexosylceramides Internal Standards (cat. no. 5040167 and 5040398, AB SCIEX). The organic phase was rated and evaporated by Savant Speedvac spd111v (Thermo Fisher Scientific) at RT for 1-2 hours. The remaining lipid pellets were stored in -80°C for further use.

Lipid pellets were reconstituted in 100% ethanol and ran on liquid chromatography electrospray ionization tandem mass spectrometry to identify several lipid classes. Sphingomyelin, cholesterol esters, ceramides, hexose-ceramides, and lactose-ceramides were measured in positive ion mode with a precursor scan of 184.1, 369.4, 264.4, 266.4, 264.4, and 264.4, respectively. Triglycerides, diglycerides and monoglycerides were measured in positive ion mode with a neutral loss scan for one of the fatty acyl moieties. Phosphatidylcholine, alkylphosphatidylcholine, alkenylphosphatidylcholine, lysophosphatidylcholine, phosphatidylethanolamine, alkylphosphatidylethanolamine, alkenylphosphatidylethanolamine, lyso phosphatidylethanolamine, phosphatidylglycerol, phosphatidylinositol, and phosphatidylserine were measured in negative ion mode by fatty acyl fragment ions. Lipid quantification was performed by scheduled multiple reactions monitoring, the transitions being based on the neutral losses or the typical product ions as described above.

Peak integration was performed with the MultiQuantTM software version 3.0.3. Lipid species signals were corrected for isotopic contributions (calculated with Python Molmass 2019.1.1). The total lipid amount and concentration per lipid class were determined with absolute values in nmol/mg DNA. The lipidomic dataset was analysed with the web-based application MetaboAnalyst 5.0 [51]. Missing values were addressed through imputation. Lipid species with >50% missing values were removed, and the remaining missing values were substituted by LoDs (1/5 of the smallest positive value of each variable). To prepare data for univariate and multivariate analyses, data were log10 transformed and Pareto scaling was applied (mean-centred and divided by the square root of the standard deviation of each variable). Groups were compared using principal component analysis and hierarchical clustering heatmaps (distance measure: Euclidean, clustering algorithm: ward, standardization: auto-scale features). This multivariate-based approach identified a technical outliner with very high lipid concentrations compared to all the other samples, which was removed from the analysis.

### 2.14. Transcriptomics

Total RNA was extracted from OPCs/pmOLs using the Quick-RNA Microprep Kit (Zymo research). Libraries were prepared from 500 ng using the QuantSeq 3’ mRNA-seq library prep kit (Lexogen), and pooled equimolar to 2 nM for single-read sequencing on the HiSeq4000 (Illumina) with settings 51-8-8. Quality control of the generated raw fastq sequence files was performed with FastQC v0.11.7 [2], and adapters were filtered with ea-utils fastq-mcf v1.05 [4]. Splice-aware alignment was performed with HiSat2 [22] against the human reference genome hg38 using default parameters. Reads mapping to multiple loci in the reference genome were discarded. Resulting BAM alignment files were handled with Samtools v1.5. [32]. Reads per gene were calculated with HT-seq Count v2.7.14 and count-based differential expression analysis was done with R-based Bioconductor package DESeq2 [39]. Principle component analysis was performed with regularized log transformation for clustering, and limma package was used to remove batch effects. Cut-off values of adjusted p-value <0.05 and log_2_ fold change >1 were used for the enhanced volcano plot. Gene set enrichment analysis was performed using ClusterProfiler [72] and STRING v12.0 [66].

### 2.15. Western blot

Mice were euthanized by an intraperitoneal injection of 60 mg/kg sodium pentobarbital followed by transcardial perfusion with saline, after which the brain, retina and optic nerve were dissected and snap-frozen in liquid nitrogen. Samples were homogenized in lysis buffer (150 mM NaCl, 50 mM Tris-HCl, 10 mM CaCl_2_dihydrate, 1% Triton-X 100, 0.05% BRIJ-35), supplemented with EDTA-free protease inhibitor cocktail (Roche Applied Science). Western blot was performed as previously described [31]. Briefly, samples were separated on a 4–12% Bis-Tris gel and transferred onto a PDVF membrane, followed by 2 h blocking with 15% milk powder (in TBST) and overnight incubation with rabbit anti-BIP (3183, Cell signaling, 1/1000 for retina, 1/500 for optic nerve) or mouse anti-CHOP (895, Cell signaling, 1/1000 for retina, 1/500 for optic nerve). Protein bands were visualized using horseradish peroxidase-labeled secondary antibodies and a luminol-based enhanced chemiluminescence kit. At least two Western blot experiments (i.e., two technical repeats) were performed for each of the lysates. Optical density measurements were normalized against a SWIFT Membrane total protein stain (G-Biosciences), and subsequently presented as a percentage relative to the naive condition.

OPCs/pmOLs were lysed by using M-PER buffer (Thermo Scientific™) containing PhosSTOP™ Phosphatase Inhibitor Cocktail (Roche Diagnostics) and complete™ Protease Inhibitor (Roche Diagnostics). Protein concentration was determined by using a Pierce™ BCA Protein Assay kit (Thermo Scientific™) and 20 μg protein was mixed with SDS-containing reducing sample buffer (Thermo Scientific™), followed by denaturation at 95°C for 10 min and loading on 4-20% ExpressPlus^TM^ PAGE gels (GenScript). The gel was allowed to run at 120V for 2 hours, followed by 0.2 μm nitrocellulose membrane transfer at 25V, 2.5A for 7 min using iBlot^TM^ 2 dry Blotting Transfer system (Thermo Fisher). The membrane was blocked in Tris-buffered saline with Tween-20 (TBST) (Sigma-Aldrich) containing 5% skim milk (Sigma-Aldrich) for 1 hour, incubated overnight with primary antibody (BiP, cat. 3183S dilution 1:1000 or WFS-1 cat. PA5-76065 dilution 1:2000) in 3% bovine serum albumin-TBST at 4°C. The next day, incubation with secondary antibody diluted in TBST was performed for 1 hour. SuperSignal™ West Pico PLUS Chemiluminescent Substrate (Thermo Scientific™) was used for chemiluminescence imaging (ImageQuant LAS4000, GE Healthcare). Beta-actin (ref. 8457, Cell Signaling, 1/1000) or vinculin (ref. V9131, Sigma, 1/1000) were used as a loading control and for normalization of the measured optical densities.

### 2.16. Immunohistochemistry and morphometric analyses

Mice were euthanized by an intraperitoneal injection of 60 mg/kg sodium pentobarbital, followed by transcardial perfusion with saline and 4% PFA. For retinal flatmounts, the eyes were post-fixated in 4% PFA for 1 hour, retinas were flatmounted and post-fixated for another hour in 4% PFA. For cryosections of the optic nerve, the nerves were immersed in 4% PFA for 30 minutes.

For wholemount immunostaining, wholemounts were frozen for 15 min at -80°C and incubated overnight at room temperature with primary antibodies for RNA-binding protein with multiple splicing (RBPMS) (ref. 1830-RBPMS, PhosphoSolutions, 1/250) or TH (ref. AB152, Millipore, 1/1000), followed by incubation with corresponding fluorophore-conjugated secondary antibodies. Images were made with a DM6 fluorescence microscope (Leica), and analysed using ImageJ software [42]. RBPMS+ retinal ganglion cells were counted using a validated automated counting method [27]. Tyrosine hydroxylase+ amacrine cell numbers were counted manually [50].

For cryosections of the optic nerve, tissues were cryoprotected in an ascending sucrose series and embedded in optimal cutting temperature medium (Tissue-Tek, Lab Tech) to make 12 μm thick longitudinal sections. Optic nerve cryosections were blocked with 20% pre-immune donkey serum for 45 min, followed by overnight incubation with the primary antibody for platelet-derived growth factor receptor alpha (PDGFRa) (ref. AF1062-SP, R&D systems, 1/200), OLIG2 (ref. 13999-1-AP, Proteintech, 1/400), or CC1 (anti-adenomatous polyposis coli clone CC1, ref. OP80, Calbiochem, 1/200) and a 2-hour incubation with the secondary fluorophore-conjugated antibody. Nuclei were counterstained with DAPI and sections were mounted with mowiol. Images were taken with an epifluorescence microscope (DM6 B, Leica Microsystems) and 20X objective. Two or three representative sections were chosen for each optic nerve, on which cells in 3 regions of interest (ROI) of 0.08 mm² were manually counted using the cell-counter plugin of ImageJ: one ROI close to the ONH, one ROI in the middle of the optic nerve and one ROI close to the optic chiasm.

For immunocytochemistry of OPCs/pmOLs, 30000 cells per well were plated in a 96-well plate in OMM. The cells were maintained for 2 days, followed by fixation with 4% PFA for 15 min at room temperature. The cells were washed, blocked and permeabilized with 5% goat serum (Dako) and 0.1% Triton X-100 (Sigma) for 1 hour. Overnight incubation with the primary antibody (Chicken anti-MBP, cat. ab9348 dilution 1:75 and Mouse anti-O4, cat. MAB1326 dilution 1:1000) diluted in 5% goat serum at 4°C was followed by secondary antibody incubation, diluted in Dako REAL Antibody Diluent (Dako) for 1 hour. Hoechst33342 (Sigma, dilution 1/2 000 in Dako REAL Antibody Diluent) was applied for nuclear counterstaining. Fluorescence imaging was performed using the Operetta High Content Imaging System (PerkinElmer) and analysis was done using automated segmenting and counting of objects using Columbus software (PerkinElmer). Briefly, the nuclear staining was segmented and used as a reference to have object count and cytoplasm segregation. The cell population was selected based on the mean fluorescence intensity in both cell regions (nuclear and cytoplasmic), using a threshold to select the positive population. The integrated fluorescence intensity was measured by dividing the mean fluorescence intensity by the cytoplasm area (in µm^2^).

### 2.17. Transmission electron microscopy and morphometric analyses

Mice were euthanized by an intraperitoneal injection of 60 mg/kg sodium pentobarbital (Dolethal, Vetoquinol) followed by transcardial perfusion with 2.4% glutaraldehyde and 4% PFA in 0.1M Na-cacodylate buffer. Optic nerves were gently dissected from the brain, while being immersed in fixative, followed by overnight post-fixation at 4°C. Next, ascending concentrations of acetone were used to dehydrate the samples, after which they were placed in a 1:1 mixture of araldite epoxy resin and acetone overnight, and embedded in araldite expoxy resin. Next, 70 nm cross-sections of the optic nerves were made, transferred to 0.7% formvar-coated copper grids and contrasted with 0.5% uranyl acetate and lead citrate. Images were made with an EM208 S electron microscope (Philips), Morada soft imaging system camera and iTEM FEI software (Olympus). ImageJ software was used for morphometric analysis of at least 100 axons (on 4 different sections) per optic nerve. Axon area and axon-plus-myelin area were measured by encircling these areas, from which inner and out axon diameter, respectively, were calculated, as well as the g-ratio. Myelin area was calculated by subtracting the axon area from the axon-plus-myelin area. White space or interaxonal space was calculated by subtracting the sum of the axon-plus-myelin areas within a given region of interest from the total area of that region. Both measures are shown relative to the total region of interest (in %). Axon density was defined by counting the number of axons (Cell Counter plugin) in a 20×16 μm region of interest.

### 2.18. Statistical analysis

The employed statistical analyses and number of mice (N) or number of experiments performed (n) are stipulated in the respective figure legends. For experiments with OPCs/pmOLs, a minimum of three independent repeats, based on at least three different differentiation batches, was performed. Statistical analyses were performed using Prism v.8.2.1 (GraphPad). Differences were considered statistically significant for two-sided p-values < 0.05 (*p < 0.05; **p < 0.01; ***p < 0.001; ****p < 0.0001).

## 3. Results

### 3.1. Progressive vision loss and neuronal dysfunction in *Wfs1* knockout mice

Vision loss due to optic neuropathy is one of the main features of WS [69]. We therefore started with a comprehensive evaluation of visual function in the *Wfs1* KO mice. *Wfs1* KO mice indeed present with visual defects early in life: contrast sensitivity is reduced from the age of 3 months (Figure 1a) and visual acuity declines starting at 6 months of age (Figure 1b). Furthermore, electroretinograms showed that the amplitude of the positive scotopic threshold response progressively declined from 3 till 7.5 months of age in *Wfs1* KO mice (Figure 1c and Figure S2), suggesting retinal ganglion cell dysfunction. A- and b-wave measurements were normal (Figure S2), indicating that specifically the retinal ganglion cells are primarily affected in the *Wfs1* KO mice. Next, visual evoked potential recordings also pointed to synaptic dysfunction and/or abnormal action potential conduction in the optic nerve, as evident from an age-dependent increase in the latency time of the visual evoked potentials (Figure 1d and Figure S2). To directly assess axonal conduction properties of retinal ganglion cell axons, *ex vivo* compound action potential (CAP) recordings in the optic nerve were performed at the oldest age (7.5 months). No differences in the CAP peak latencies between the genotypes were observed (Figure 1e-f), suggesting that action potential conduction speed is not overtly perturbed. However, when optic nerves were challenged with increasing stimulation frequencies, a more pronounced CAP peak decline at high frequencies was visible in *Wfs1* KO mice compared to WT littermates (Figure 1g), suggesting an increase in conduction blocks when axons are required to fire at high rates. Notably, the post-stimulation CAP peak recovery kinetics appeared normal in *Wfs1* KO mice, implying that oligodendroglial potassium clearance is not overtly affected [37]. Finally, *in vivo* imaging of axonal transport in the optic nerve via manganese-enhanced MRI, showed an optic nerve diameter that tends to be smaller (with a significant difference at 6 months), and reduced innervation of the superior colliculus –*i.e.* reduction in the Mn^2+^-labeled area of the superior colliculus after an intravitreal injection with Mn^2+^– already at 3 months of age (Figure 1h-i). Altogether, compared to WT littermates, *Wfs1* KO mice display an age-dependent deterioration of visual function, which culminates into progressive retinal ganglion cell dysfunction, higher susceptibility to activity-induced conduction blocks, and vision loss.

**Figure 1.**
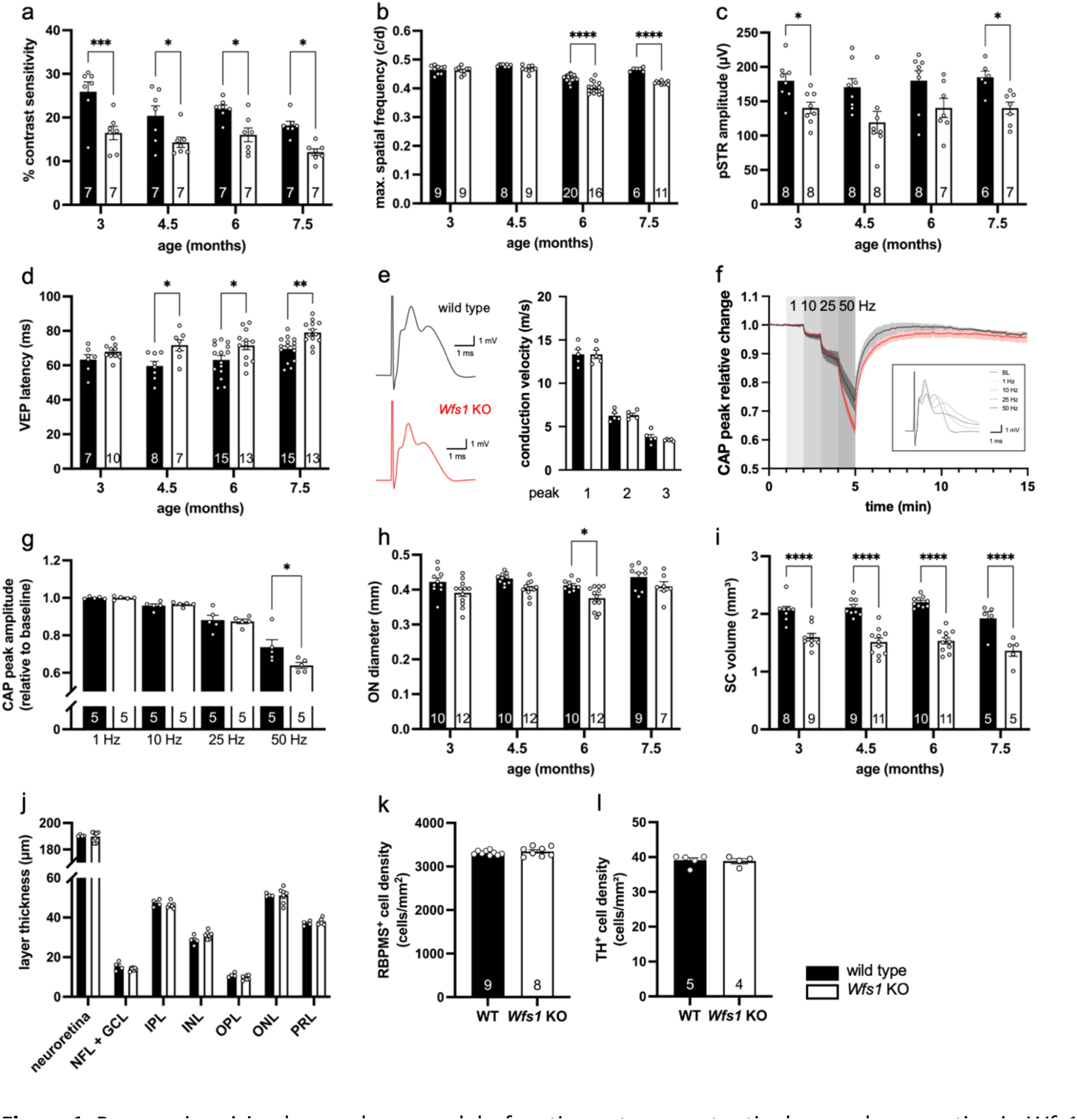
Progressive vision loss and neuronal dysfunction yet no overt retinal neurodegeneration in *Wfs1* knockout mice. **a, b** Visual function is affected in *Wfs1* KO compared to WT mice, with contrast sensitivity (**a**) being affected earlier in life than visual acuity (**b**). Two-way ANOVA with Sidak’s multiple comparisons; F_1,80_=65.04 and p<0.0001 for genotype, F_3,80_=90.47 and p<0.0001 for age, for contrast sensitivity; F_1,48_=40.33 and p<0.0001 for genotype, F_3,48_=5.524 and p=0.0024 for age, for visual acuity. **c** The amplitude of the positive scotopic threshold response is reduced in *Wfs1* KO mice. Mixed-effects analysis with Sidak’s multiple comparisons; F_1,15_=18.46 and p=0.006 for genotype, F_3,31_=1.006 for age. **d** Latency of visual evoked potentials increases with age in *Wfs1* KO compared to WT mice. Two-way ANOVA with Sidak’s multiple comparisons; F_1,80_=24.90 and p<0.0001 for genotype, F_3,80_=6.720 and p=0.0004 for age. **e** Representative CAP traces for wild type and *Wfs1* KO mice and the conduction velocity of the individual peaks at baseline (0.4 Hz). **f, g** Relative changes in peak amplitude, in response to increasing stimulation frequencies, show a larger CAP peak amplitude (P2) drop at 50 Hz in *Wfs1* KO nerves compared to WT (**g**). Unpaired t-test; t_8_=2.317 and p=0.0492. **h-i** Mn^2+-^enhanced MRI of the retinocollicular projection reveals a reduced optic nerve diameter **(h**) and superior colliculus innervation **(i**). Two-way ANOVA with Sidak’s multiple comparisons; F_1,74_=21.41 and p<0.0001 for genotype, F_3,74_=3.588 and p=0.0178 for age, for optic nerve; F_1,60_=142.6 and p<0.0001 for genotype, F_3,60_=3.331 and p=0.0253 for age, for superior colliculus. **j** Analysis of the thickness of the retinal layers, as measured by optical coherence tomography, shows no differences between *Wfs1* KO and WT mice at 7.5 months of age. Multiple unpaired t-tests: t_9_=0,3632 for neuroretina, t_9_=1,587 for NFL + GCL, t_9_=1,137 for IPL, t_9_=1,962 for INL, t_9_=1,932 for OPL, t_9_=0,07964 for ONL, t_9_=0,9323 for PRL. **k, l** Cell counts of retinal ganglion cells (**k**) and dopaminergic amacrine cells (**l**) reveal similar numbers in 7.5-month-old *Wfs1* KO and WT animals. Unpaired t-test; t_15_=20.8088 for ganglion cells; t_7_=0.2544 for amacrine cells. Data presented as mean ± SEM, number of animals (N) depicted in the bar graphs.

Analysis of the structural integrity of the retina, via *in vivo* optical coherence tomography revealed no apparent thinning nor swelling of the retinal layers in *Wfs1* KO *versus* WT mice (Figure 1j). Immunohistological stainings at 7.5 months of age confirmed that there were no differences in the number of retinal ganglion cells (Figure 1k) or dopaminergic amacrine cells (Figure 1l) in the *Wfs1* KO mice compared to their WT littermates. Thus, despite the large body of evidence for neuronal dysfunction, we did not observe any signs of neurodegeneration in the retina of *Wfs1* KO mice up till the age of 7.5 months.

### 3.2. ER stress as potential underlying mechanisms of disease

As a potential underlying pathological mechanism of WS, we assessed ER stress. Western blotting showed increased levels of immunoglobulin heavy chain binding protein (BiP) –correlating with the accumulation of unfolded proteins in the ER lumen– (Figure 2c-d) yet not of C/EBP homologous protein (CHOP) –usually observed in case of severe/unresolved ER stress– (Figure 2e-f), in both retina and optic nerve lysates. Notably, the optic nerve was affected sooner (already at 3 months) than the retina (6 months).

**Figure 2.**
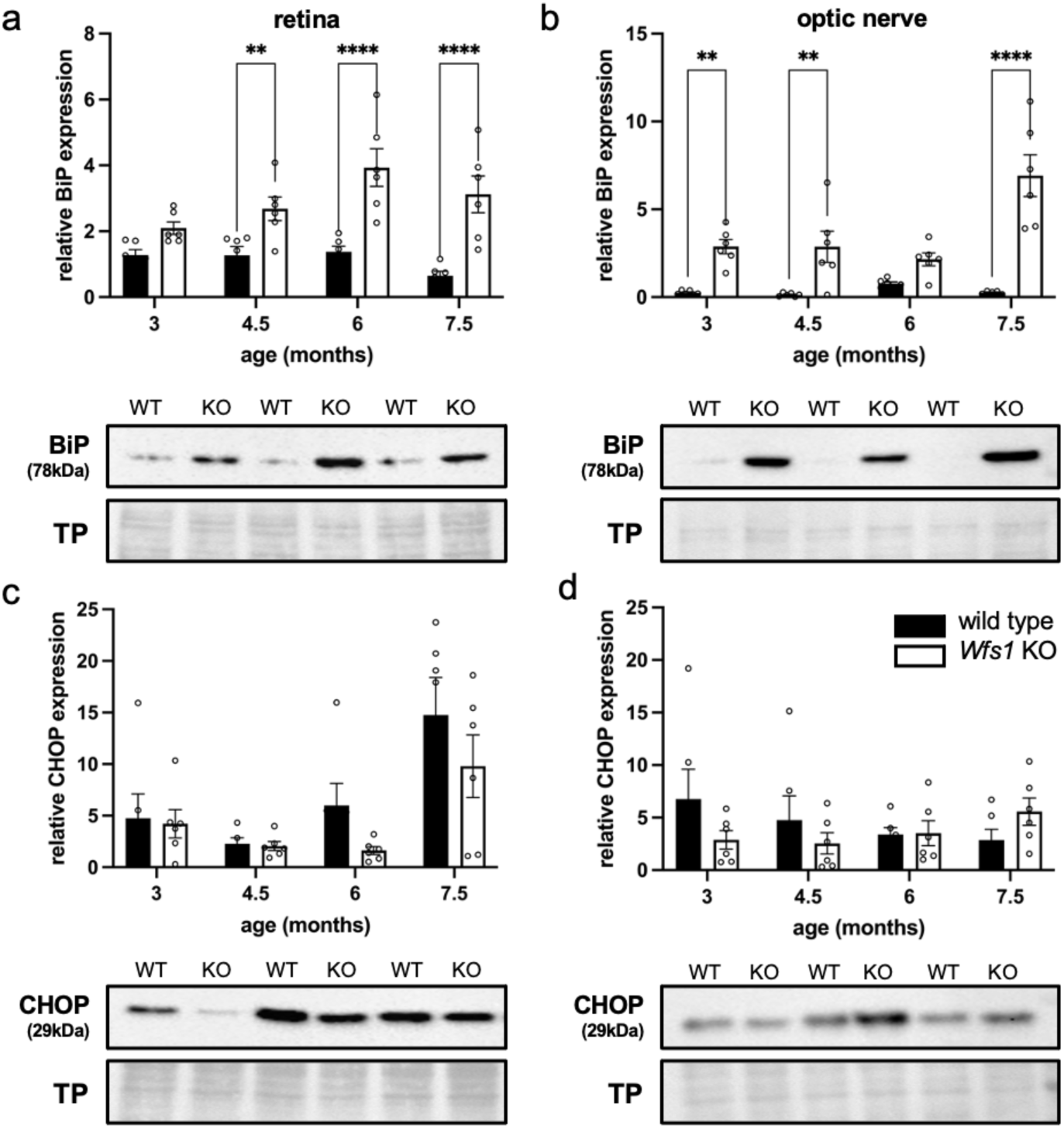
Activation of the unfolded protein response but no unresolved ER stress in the retina and optic nerve of *Wfs1* KO mice. **a, b** Elevated BiP levels indicate that there is accumulation of unfolded proteins in the ER lumen in the retina (**a**) and optic nerve (**b**) of *Wfs1 KO* mice, but not of WT controls; starting from the age of 4.5 and 3 months, respectively. Two-way ANOVA with Sidak’s multiple comparisons; F_1,40_=55.52 and p<0.0001 for genotype, F_3,40_=2.992 and p=0.0421 for age, with p=0.0997 for 3 months, p=0.0061 for 5 months, p<0.0001 for 6 months, p<0.0001 for 7.5 months, for retina; F_1,39_=67.94 and p<0.0001 for genotype, F_3,39_=6.239 and p=0.0015 for age, with p=0.0022 for 3 months, p=0.0015 for 4.5 months, p=0.0956 for 6 months, p<0.0001 for 7.5 months, for optic nerve. **c, d** CHOP expression levels are similar for both genotypes in the retina (**c**) and optic nerve (**d**). Two-way ANOVA with Sidak’s multiple comparisons; F_1,39_=2.763 for genotype, F_1,19_=8.846 and p=0.0036 for age, for retina; F_1,10_=0.5544 for genotype, F_3,30_=0.3121 for age, for optic nerve. Data presented as mean ± SEM, N=6. TP: total protein stain.

### 3.3. Detailed analysis of the optic nerve suggests minor oligodendroglia defects

The central research question of this study is whether oligodendroglia may be the catalyzers of the neurodegeneration observed in WS. The observed increase in latency of the visual evoked potential recordings and the altered axonal conduction strength may be associated with myelin deficits. Hence, next, we further exploited the optic nerve to characterize in detail the oligodendroglia and myelin integrity in *Wfs1* KO *versus* WT mice. First, analysis of immunostainings on longitudinal optic nerve cryosections revealed equal numbers of OLIG2^+^ oligodendroglial cells (i.e., OPCs and oligodendrocytes) (Figure 3b) and CC1^+^ mature oligodendrocytes at 7.5 months of age (Figure 3c). However, cell counting of PDGFRα^+^ OPCs did show a reduced number of these cells in the optic nerve of *Wfs1* KO mice compared to WT controls (Figure 3d). Analyses at younger ages indicate that this is not a developmental defect, but that this depletion arises around the age of 6 months (Figure 3d). Next, we determined the g-ratio of optic nerve axons on TEM images and found a higher g-ratio in the *Wfs1* KO mice –i.e., myelin thinning– at 7.5 months of age (Figure 3e). This defect was subtle, as the average myelin area was similar for both genotypes (Figure 3f). Furthermore, at this age, axon density, axon diameter and the empty space in between axons were identical in *Wfs1* KO and WT mice (Figure 3g-i). Together, these findings suggest a complex disease manifestation, with only late and subtle thinning of the myelin sheath and a reduction of the OPC pool.

**Figure 3.**
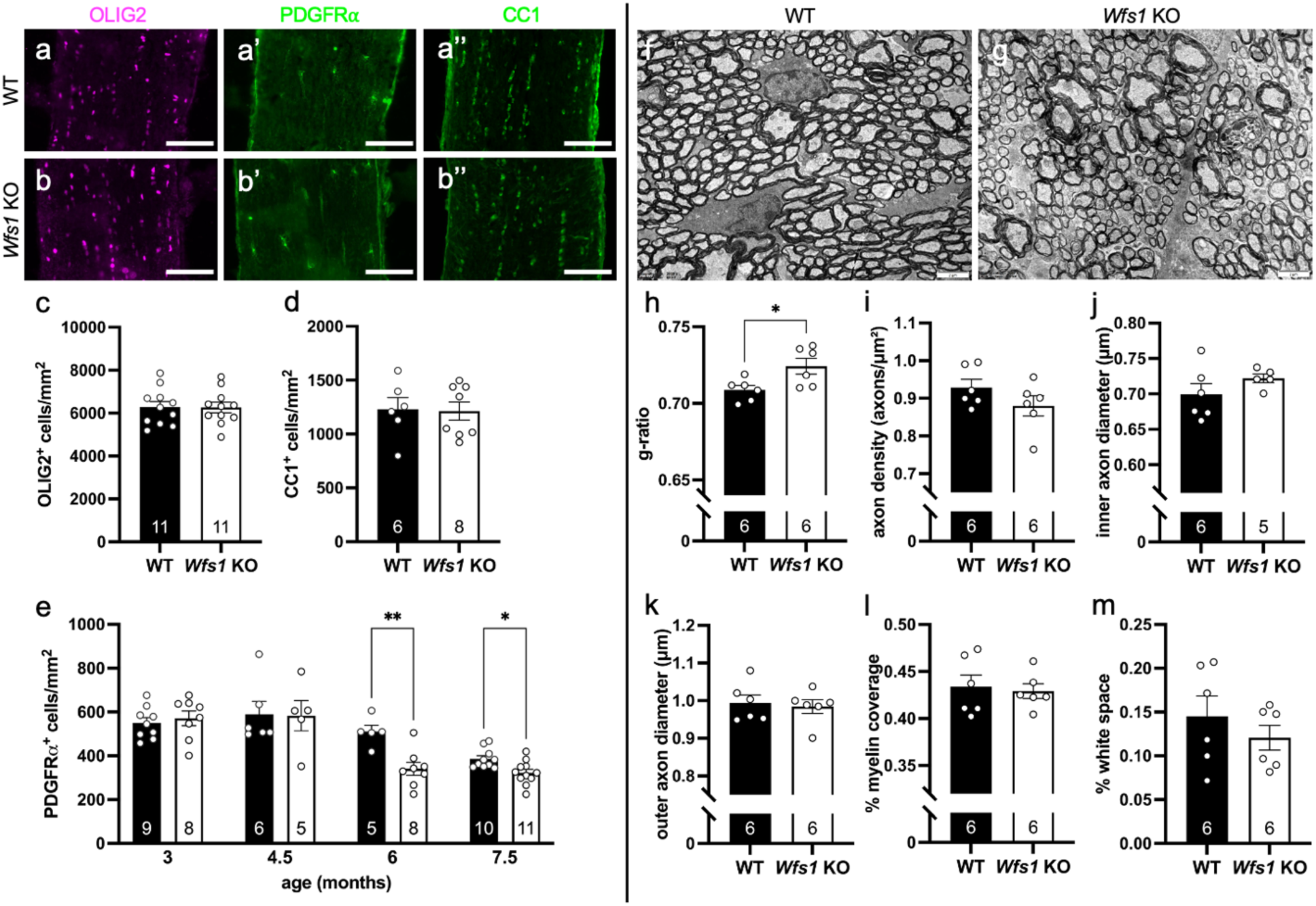
A reduction in the number of OPCs and minor myelin defects in the *Wfs1* KO optic nerve. **a-e** Immunohistological stainings with different oligodendroglia lineage markers of WT (**a**) and *Wfs1* KO optic nerves (**b**) show equal cell numbers of OLIG2^+^ oligodendroglial lineage cells (unpaired t-test, t_20_=0.05208) (**c**), and CC1^+^ mature oligodendrocytes (unpaired t-test, t_12_=0.1220) (**d**); but a reduced number of PDGFRα^+^ OPCs in *Wfs1* KO optic nerves compared to WT at 6 and 7.5 months of age (**e**). Mixed-effects analysis with Sidak’s multiple comparisons; F_1,54_=5.512 and p=0.0226 for genotype, F_2,27_=25.49 and p<0.0001 for age; with p=0.0092 for WT *versus* Wfs1 KO at 6 months and p=0.0317 for WT *versus* Wfs1 KO at 7.5 months. **f-m** Transmission electron microscopy study of the optic nerve of 7.5-month-old WT (**f**) and *Wfs1* KO (**g**) mice reveals a difference between genotype for g-ratio (unpaired t-test, t_2.6_=10, p=0.0258) (**h**), but not for axon density (unpaired t-test, t_1.4_=10) (**i**), inner axon diameter (unpaired t-test, t_1.3_=9) (**j**), outer axon diameter (unpaired t-test, t_0.3_=10) (**k**), myelin area (unpaired t-test, t_0.3_=10) (**l**) or empty space in between axons (unpaired t-test, t_0.9_=10) (**m**). Scale bar 100 μm (a-b), 2 μm (f-g). Data presented as mean ± SEM, number of animals (N) depicted in the bar graphs.

### 3.4. The eye as a window to the brain

Although only minor morphological changes were apparent in the retina and optic nerve, structural neurodegeneration was detected in the brain of *Wfs1* KO mice via MRI scans. In addition to shrinkage of superior colliculus volume and optic nerve diameter, as observed via Mn^2+^-enhanced MRI (Figure 1g-h), we observed more manifestations of neurodegeneration. First, MRI showed an overall reduction in brain volume (Figure 4a), which likely correlates with the overall smaller posture of the *Wfs1* KO mice. Specific brain structures that were smaller in *Wfs1* KO mice compared to WT controls, after correction for smaller brain volume, were the brain stem (starting from 3 months) (Figure 4b), cerebellum (starting from 4.5 months) (Figure 4c) and superior colliculus (starting from 3 months) (Figure 4d), but not olfactory bulb (Figure 4e) nor cortex (Figure 4f). Second, analysis of fractional anisotropy at the age of 7.5 months revealed significantly reduced values, indicative of reduced myelin integrity, in the medulla, pons, cerebellum and superior colliculus, but not in the corpus callosum, visual, motor or auditory cortex (Figure 4g). Mean diffusivity values were only different in the cerebellum (Figure 4h). In conclusion, these findings mirror the results from MRI studies performed in WS patients (reviewed in [63]) and point out that WS manifests as a disease affecting both grey and white matter of the brain stem, cerebellum and retinocollicular pathway.

**Figure 4.**
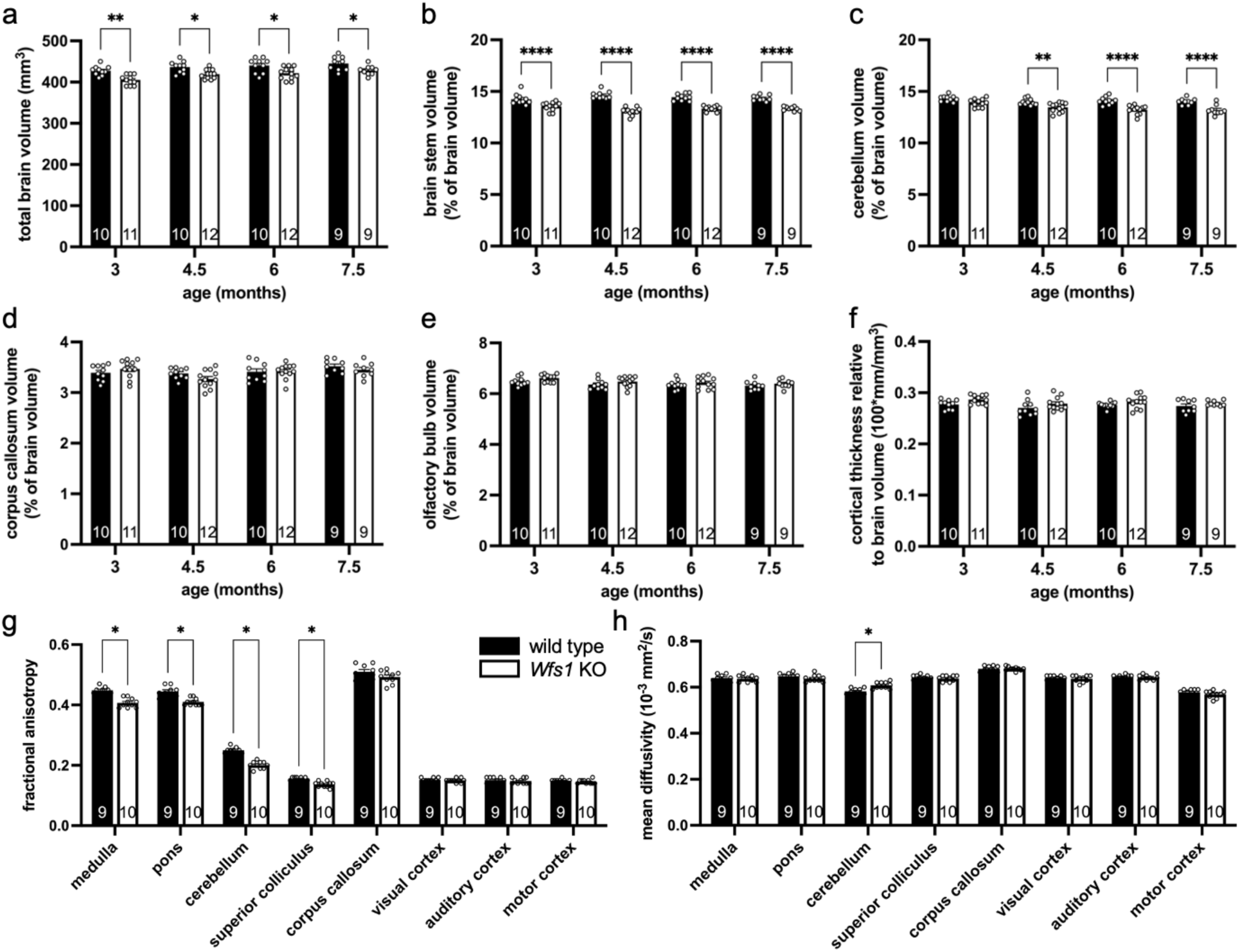
White and grey matter loss in the brain of *Wfs1* KO mice. Structural MRI and diffusion tensor imaging study of *Wfs1* KO *versus* WT mice from age 3 till 7.5 months. **a** Total brain volume is reduced in *Wfs1* KO mice of all ages. Two-way ANOVA with Sidak’s multiple comparisons; F_1,75_=37.23 and p<0.0001 for genotype, F_3,75_=7.709 and p=0.0001 for age, with p=0.0015 for 3 months, p=0.0217 for 4.5 months, p=0.0108 for 6 months, p=0.0490 for 7.5 months. **b** Brain stem volume of *Wfs1* KO animals is also reduced at all ages. Two-way ANOVA with Sidak’s multiple comparisons; F_1,75_=203.2 and p<0.0001 for genotype, F_3,75_=0.09256 for age, with p<0.0001 *Wfs1* KO *versus* WT at all ages. **c** The volume of the cerebellum is smaller in *Wfs1* KO mice starting from 4.5 months. Two-way ANOVA with Sidak’s multiple comparisons; F_1,75_=59.87 and p<0.0001 for genotype, F_3,75_=6.754 and p=0.0004 for age, with p=0.0570 for 3 months, p=0.0085 for 4.5 months, and p<0.0001 for 6 and 7.5 months. **d, f** The volume of the corpus callosum (**d**), olfactory bulb (**e**), and the thickness of the cortex (**f**) is similar between genotypes. Two-way ANOVA with Sidak’s multiple comparisons; F_1,75_=0.5229 for genotype, F_3,75_=3.910 and p=0.0119 for age, for corpus callosum; F_1,75_=6.116 and p=0.0157 for genotype, F_3,75_=4.160 and p=0.0088 for age, for olfactory bulb; F_1,75_=9.389 and p=0.0030 for genotype, F_3,75_=1.812 for age, for cortical thickness. **g** Fractional anisotropy, studied in 7.5-month-old *Wfs1* KO and WT mice, is different between genotypes for medulla (t_17_=6.753, p=0.000024), pons (t_17_=5.000, p=0.000548), cerebellum (t_17_=9.618, p<0.000001) and superior colliculus (t_17_=5.183, p=0.000449), but not for corpus callosum (t_17_=1.705), visual cortex (t_17_=0.7255), auditory cortex (t_17_=0.7334) nor motor cortex (t_17_=0.9602). Multiple unpaired t-tests. **h** Mean diffusivity studied in 7.5-month-old *Wfs1* KO and WT mice, is different between genotypes only for cerebellum (t_17_=4.144, p<0.005416). Multiple unpaired t-tests. Data presented as mean ± SEM, number of animals (N) depicted in the bar graphs.

### 3.5. Transcriptomics study of WS patient iPSC-derived oligodendrocytes

To generate more mechanistic insights into the contribution of oligodendroglia to WS pathology, we next investigated the function of these cells using WS patient-derived iPSCs as well as their isogenic controls that were differentiated into oligodendroglia. Creation of these engineered iPSC lines is described in supplement, and quality controls and characterization are shown in figures S3 to S5. First, we confirmed that oligodendroglia can be generated from *WFS1* mutant iPSCs by *SOX10* overexpression. Over the 24 days differentiation, we detected increasing transcript levels of *OLIG1*, *OLIG2* and *NKX2.2* lineage markers in *WFS1* mutant and isogenic iPSC progeny, and with >80% of the mutant and isogenic cells staining positive for O4 on DIV24 (Figure 5a-f). A small proportion of these cells (∼10%) also expressed MBP (Figure 5c) and culturing these oligodendroglia on electrospin fibers revealed MBP^+^ extensions on the fibers. Hence, we defined the day 24 iPSC progeny as a mixed population of OPCs and pmOLs.

**Figure 5.**
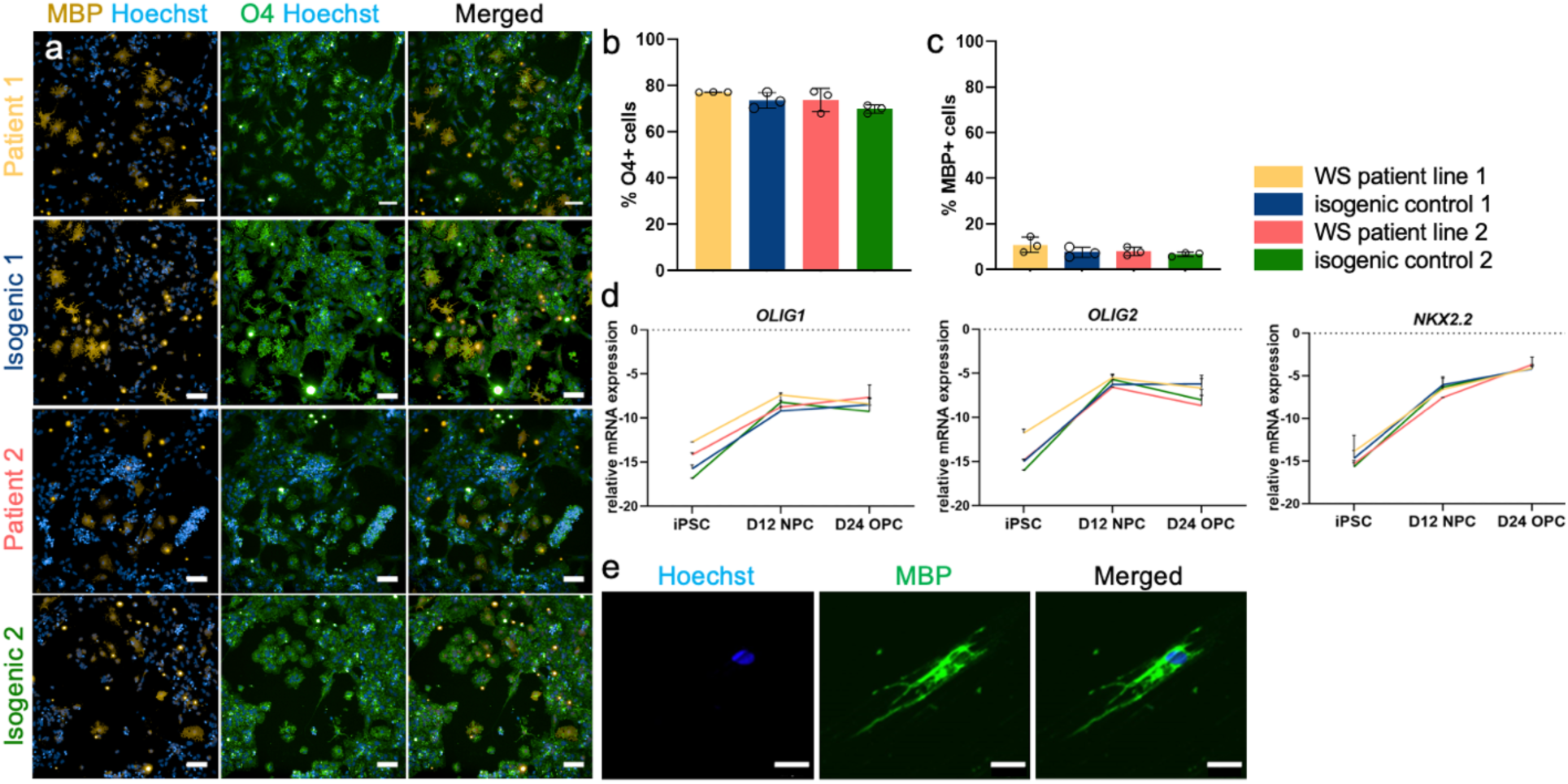
Presence of WFS1 mutation does not affect iPSC-derived OPCs/pmOLs differentiation. **a** Immunocytochemistry for O4 and MBP on OPCs/pmOLs generated from WS patient iPSC lines 1 and 2, with their respective isogenic controls. Scale bar: 100 μm. **b**-**c** Quantification of the number of immunostained O4^+^ (**b**) and MBP^+^ cells (**c**) present at DIV24. **d** RT-qPCR for oligodendroglia lineage markers *OLIG1*, *OLIG2*, and *NKX2.2* throughout the OPC/pmOL differentiation of WS patient and isogenic lines. **e** Immunostaining for MBP on DIV24 oligodendroglia (23 days in vitro, DIV24) cultured on electrospinning fibers for two weeks. Scale bar: 25 μm. Unpaired t-test; t_4_=1.840 for WS patient 1 vs. isogenic 1 and t_4_= 1.268 for WS patient 2 vs. isogenic 2, for O4; t_4_=1.406 for WS patient 1 vs. isogenic 1 and t_4_=1.062 for WS patient 2 vs. isogenic 2, for MBP. Data presented as mean ± SEM, n= 3 (a,c,d), n=1 (e) and n=3 (f).

Following confirmation that WFS1 expression levels were restored in both *WSF1* isogenic iPSCs [47] and OPCs/pmOLs (Figure S6), we performed a comparative transcriptomics study of the two pairs of WS patient iPSC-derived *versus* their isogenic control OPCs/pmOLs. A principal component analysis showed no clear clustering of patient line 1 and its isogenic control (Figure 6a), and only 8 differentially expressed genes (adjusted p-value<0.05) were found. These were mitochondrial genes (*MT-TA, MT-TF, MT-TH, MT-TK, MT-TM, MT-TN*) and *WFS1*, which were downregulated in the mutant cells, and *ZNF677,* of which the expression was upregulated (Figure 6b). For patient line 2, clustering of the mutant *versus* isogenic control cells was observed (Figure 6c) with a total of 895 differential transcripts (adjusted p-value >0.05) (Figure 6d). In both lines, oligodendrocyte lineage (*OLIG1, OLIG2, NKX2.2*) and differentiation (*RLBP1, PDGFRA, CSPG4, GPR17, CMIP2, PNPLA3, HAPLN2, APOD*) markers were not significantly altered in mutant *versus* isogenic OPCs/pmOLs, and, besides reduced *OMG* transcript levels in patient line 2, also expression of myelin components *MBP, MAG, PLP1, MOG* and *CNP* was similar. Furthermore, looking at transcripts with a link to known WS disease processes, in patient line 2, ER membrane was found twice in the top-5 cellular component GO terms, with genes connected to intracellular Ca^2+^ homeostasis and ER functions (*AKAP6, RYR2, CAMK2B, JPH4*). *RTN1*, encoding an ER-associated protein involved in neuroendocrine secretion and considered a specific marker for neurological disease, was found to be one of the most downregulated genes (Figure 6e).

**Figure 6.**
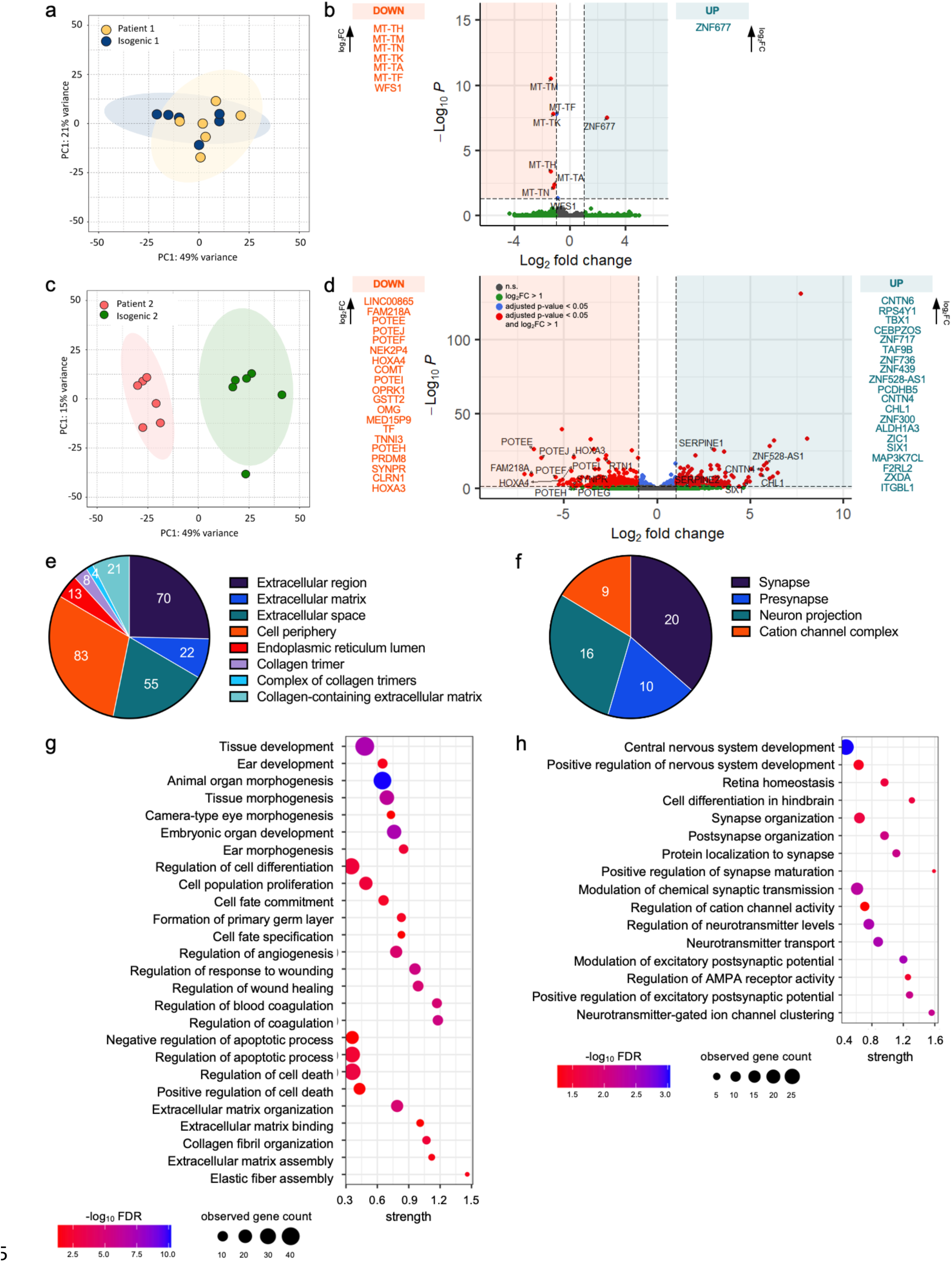
Transcriptomics analysis of WS patient iPSC-derived OPCs/pmOLs and isogenic controls. **a** Principal component analysis of patient line 1 showing no distinct clustering. **b** Enhanced volcano plot showing differentially expressed genes for *WFS1* mutant *versus* isogenic control of patient line 1. Cut-off: adjusted p-value < 0.05 and log_2_FC > 1. Side panels showing top-20 differentially expressed genes ranked based on log_2_FC (highest at the top). **c** Principal component analysis of patient line 2 showing clustering of *WFS1* mutant *versus* isogenic control samples. **d** Enhanced volcano plot showing differentially expressed genes for *WFS1* mutant *versus* isogenic control of patient line 2. Cut-off: adjusted p-value < 0.05 and log_2_FC > 1. Side panels showing top-20 differentially expressed genes ranked based on log_2_FC. **e-f** Functional enrichment analysis of differentially expressed genes in *WFS1* mutant OPCs/pmOLs from patient line 2, showing observed gene counts per GO Component for upregulated (**e**) and downregulated transcripts (**f**). **g-h** Functional enrichment analysis of differentially expressed genes in *WFS1* mutant OPCs/pmOLs from patient line 2, showing selected GO Processes associated with upregulated (**g**) and downregulated transcripts (**h**). n=6

Remarkably, as seen previously in *WFS1* mutant cells [20, 21], several Serpin family members were upregulated, most prominently paralogs *SERPINE1* and *SERPINE2* (Figure 6d). Besides their well-described role in blood coagulation, Serpins are also involved in preserving synaptic networks and their upregulation has been linked to several neurodegenerative diseases, including multiple sclerosis (MS) and amyotrophic lateral sclerosis (ALS). In line with known Serpin functions, pathways and processes connecting to morphogenesis and development, cell differentiation and cell fate commitment, as well as wound healing and blood coagulation, regulation of cell death/apoptosis, and extracellular matrix, were upregulated. Finally, in patient line 2, we observed that suppressed pathways in the mutant cells were mostly related to CNS development, synaptic signalling and regulation of neurotransmitter transport, with downregulated GO processes with a role in the cellular and molecular organisation of axo-glial synaptic junctions, and expression of the majority of the top-200 genes localising to neuronal compartments, synapse/synaptic vesicles, and cell junction (Figure 6f).

Examining the human diseases that have been linked to the differentially expressed genes in this study, we found that many of the top-20 genes correlate with neurodevelopmental or neurodegenerative WS-like syndromes, characterized by hearing and/or vision loss, or diabetes. These include, in patient line 1, rare mitochondrial non-syndromic sensorineural deafness (*MT-TA, MT-TH, MT-TK, MT-TN, MT-TS1, MT-TW*), and, in patient line 2, Baraitser-Winter syndrome (*POTEE, POTEF, POTEG, POTEH, POTEI, POTEJ*), cone-rod dystrophy 13 (*FAM218A)*, Athabaskan brainstem dysgenesis syndrome (*HOXA3* and *HOXA4)*, nutritional optic neuropathy (*SYNPR),* Cockayne syndrome A (*ZNF528-AS1)*, autosomal dominant deafness type 23 and branchio-otorenal spectrum disorder (*SIX1)*, and 3p deletion syndrome (*CNTN4* and *CHL1)*. Taken together, in both patient lines, we found little evidence of differential expression of genes related to oligodendroglia function or within the oligodendroglia lineage cell compartment. Some differentially expressed genes localize to the mitochondria or ER, and play a role in intracellular Ca^2+^ homeostasis, ER and mitochondrial function.

### 3.6. Mechanistic studies of WS disease processes in iPSC-derived oligodendroglia

Based on the results of the transcriptomics study, we next evaluated several cellular functions that have been linked to WS, and for which altered transcripts were found –mostly related to ER, in patient line 2. As delineated in the introduction, mutant WFS1 causes a typical MAM phenotype, associated with decreased ER Ca^2+^ stores, increased ER stress susceptibility, aberrant mitochondrial function, as well as defects in phosphatidylcholine generation from phosphatidylethanolamine –a lipid exchange and transport process that typically requires close contacts between the ER and mitochondria. We therefore assessed these different processes in the mutant and isogenic oligodendroglia. Accumulation of unfolded proteins and ER stress was assessed via RT-qPCR gene expression analysis of markers of the UPR pathway (*BiP*, *CHOP*, *IRE1a*, *ATF6* and *PERK*) in basal conditions as well as upon treatment with ER-stress inducers. To evoke ER stress, we opted to use thapsigargin, an irreversible inhibitor of SERCA pumps thus depleting the ER Ca^2+^ stores, and tunicamycin, an inhibitor of the N-linked glycosylation of proteins, both resulting in the accumulation of unfolded proteins in the ER lumen. As anticipated, the addition of both thapsigargin and tunicamycin elevated the expression of ER stress markers (Figure 7a), however, no significant differences were observed between mutant and isogenic control cells. Western blot analysis for BiP protein levels (with tunicamycin treatment) confirmed these data (Figure 7b). Furthermore, an XBP1 splicing assay (with thapsigargin treatment) (Figure 7c), measuring *bona fide* ER stress, also did not reveal an increased susceptibility to ER stress induction in WS patient compared to isogenic OPCs/pmOLs. Second, real-time cell metabolic analysis (Seahorse XF24, Agilent) disclosed unaltered mitochondrial function, with basal respiration, maximal respiration, ATP production, spare respiratory capacity and extracellular acidification rate unaltered in both pairs of mutant and isogenic oligodendroglia (Figure 7d). Third, to assess ER-mitochondria interactions, we performed a quantitative analysis of MAMs with a proximity ligation assay using antibodies against vesicle-associated membrane protein-associated protein B (VAPB), on the ER membrane, and protein tyrosine phosphatase interacting protein 51 (PTPIP51), on the outer mitochondrial membrane, as previously used to identify a MAM phenotype in an iPSC model of ALS [76]. However, here, we did not identify any differences between mutant and isogenic iPSC-oligodendroglia (Figure 7e-f). Normal functionality of the ER was further confirmed via intracellular Ca^2+^ measurements. ATP and acetylcholine were used to stimulate cell-surface receptors and thereby IP_3_R-mediated Ca^2+^ release, which was detected by the Cal-520 dye as a Ca^2+^ sensor. No difference in terms of area under the curve and peak amplitudes (F_peak_-F_baseline_) was observed between the control and WS patient OPCs/pmOLs, thus indicating preserved Ca^2+^ storage in the ER (Figure 8 and Figure S7). Finally, further confirming a preserved MAM phenotype, lipidomic profiling showed that no lipid classes were significantly different in WS *versus* isogenic oligodendroglia, including phosphatidylcholine, phosphatidylethanolamine and phosphatidylinositol (Figure S8) –which are among the phospholipids present in the mitochondrial membrane and typically affected in case of a loss of MAMs. In summary, these experiments lead us to conclude that OPCs/pmOLs derived from WS patient iPSCs do not display any signs of ER abnormalities or chronic ER stress, mitochondrial dysfunction, or abnormal ER-mitochondria interactions.

**Figure 7.**
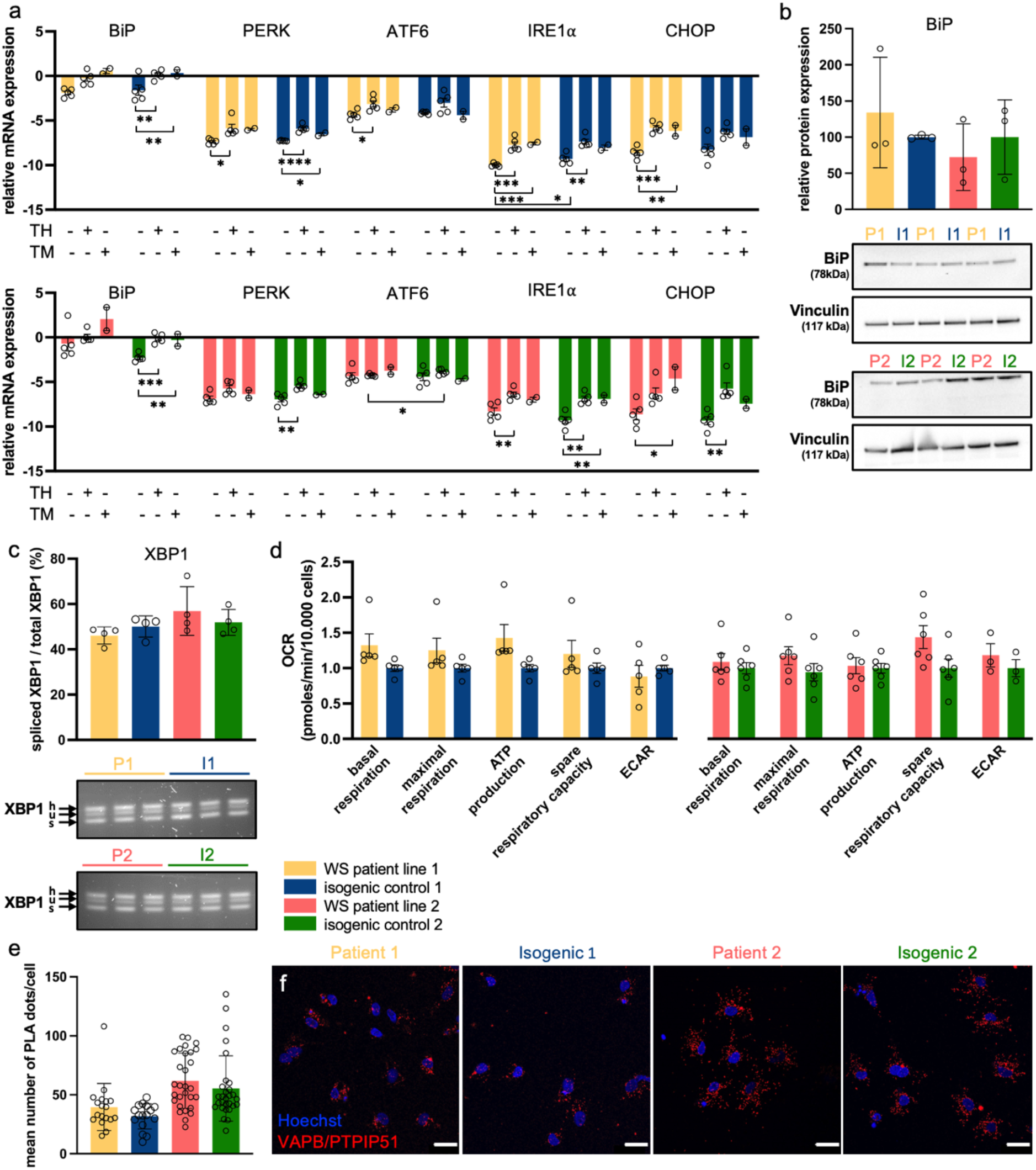
Mechanistic studies of WS disease processes in iPSC-derived OPCs/pmOLs. **a** RT-qPCR gene expression analysis of ER stress markers *BiP*, *PERK*, *ATF6*, *IRE1*α and *CHOP* in WS patient line 1 and 2, and their respective isogenic control OPCs/pmOLs, with/without ER stressors thapsigargin (TH) (2μM, 3 hours) or tunicamycin (TM) (1μg/ml, 24 hours). **b** Protein expression levels of BiP after TM treatment in WS patient OPCs/pmOLs and isogenic controls (% relative to isogenic control). Unpaired t-test; t_4_=0.7714 for WS patient 1 vs. isogenic 1 and t_4_=0.6923 for WS patient 2 vs. isogenic 2. **c** Ratio of spliced to total XBP1 mRNA in WS patient OPCs and isogenic controls after TH treatment (2μM, 3 hours). h: hybrid, s: spliced, u: unspliced XBP1 mRNA. Unpaired t-test; t_6_=1.324 for WS patient 1 vs. isogenic 1 and t_6_=0.8214 for WS patient 2 vs. isogenic 2. **d** Bioenergetic profile of mutant and isogenic control OPCs/pmOLs after addition of mitochondrial stressors, oligomycin, FCCP and rotenone. A Seahorse mitostress assay was performed to measure basal respiration, maximal respiration, ATP production, spare respiratory capacity, and extracellular acidification rate (ECAR). **e-f** Proximity ligation assay to assess proximity between ER and mitochondria in WS patient OPCs/pmOLs and isogenic controls. Unpaired t-test; t_32_=1.401 for WS patient 1 vs. isogenic 1 and t_56_=0.9760 for WS patient 2 vs. isogenic 2. Data presented as mean ± SEM. n=5 for non-treated and TH, and n=2 for TM (a), n=3 (b), n=4 (c), n=5 for patient line 1 and n=3-6 for patient line 2 (d), n=3 with at least 16 images and n>100 nuclei/condition (e). Scale bar 25 μm (f). Statistics for panel a and d are reported in supplementary table S2. P1: WS patient line 1, I1: isogenic line 1, P2: WS patient line 2, I2: isogenic line 2.

**Figure 8.**
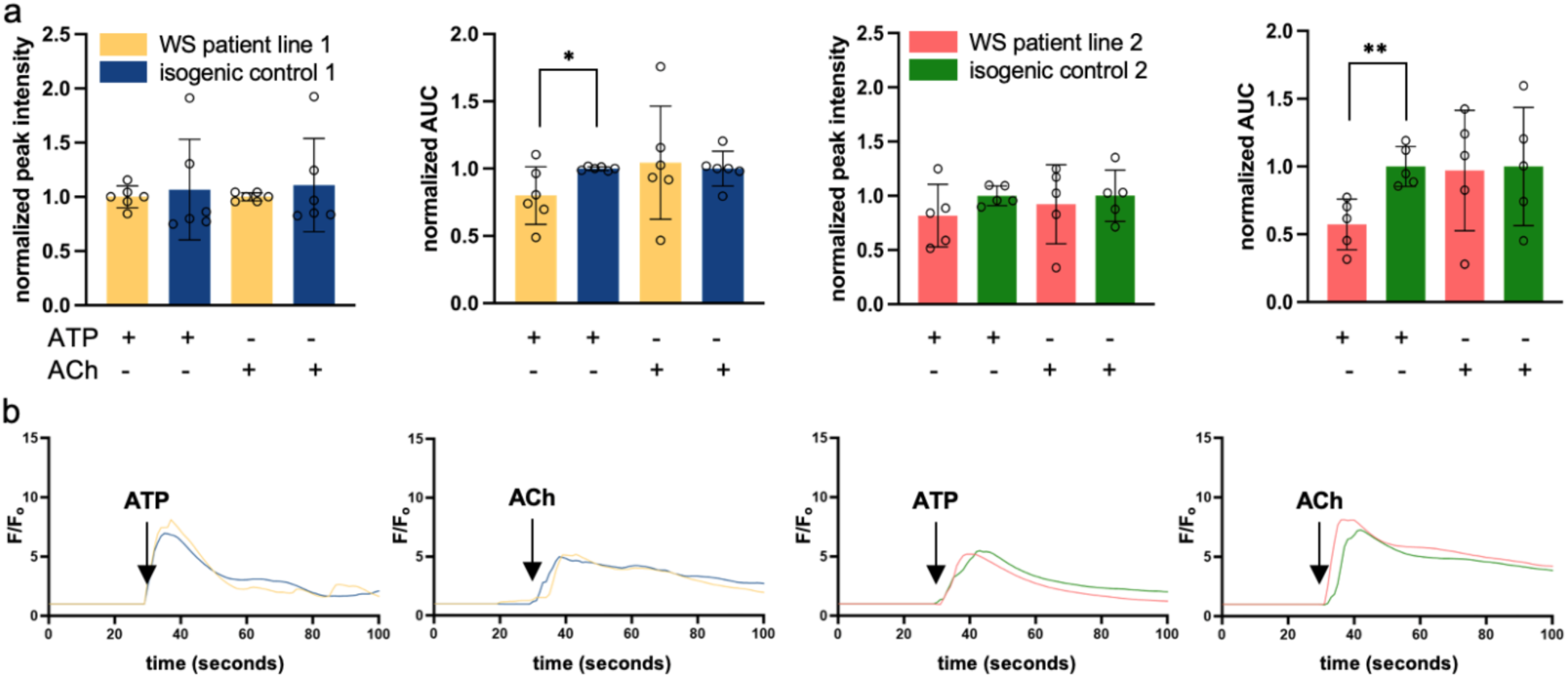
Mechanistic studies of WS disease processes in iPSC-derived OPCs/pmOLs (continued). **a** Quantification of cytosolic Ca^2+^ release by WS patient and isogenic OPCs/pmOLs after pharmacological activation of IP_3_R-mediated Ca^2+^ release by addition of 10 μM ATP or 10 μM acetylcholine (ACh) to the culture medium. **b** Average traces of cytosolic Ca^2+^ release by WS patient and isogenic OPCs/pmOLs after pharmacological activation of IP_3_R-mediated Ca^2+^ release. Data presented as mean ± SEM. n=6 for patient line 1 and n=5 for patient line 2. Statistics are reported in supplementary table S3.

## 4. Discussion

WS typically manifests as a combination of early-onset diabetes mellitus, progressive optic nerve atrophy, diabetes insipidus and sensorineural hearing loss. However, it is also associated with a number of neurological complications and psychiatric manifestations. Several studies have proven ER stress-mediated death of beta cells in WS, and therapeutic potential for diabetes medications such as glucagon-like peptide 1 receptor agonists [17, 44, 45]. In contrast, the disease mechanisms underlying the neurological and psychiatric manifestations of WS are less well understood, and recently oligodendrocytes have gotten increased attention. Indeed, similar to beta cells, by producing enormous amounts of plasma membrane during the myelination process, oligodendrocytes may be particularly susceptible to disruptions of the secretory pathway and ER stress [33]. Furthermore, beside myelinating axons, oligodendrocytes also support neuronal function and survival via metabolic coupling [37, 48]. ER stress in myelinating cells and loss of oligodendrocyte metabolic support have been shown to contribute to the pathogenesis of various demyelinating disorders, including MS, Charcot-Marie-Tooth disease, Pelizaeus-Merzbacher disease, vanishing white matter disease and ALS [33]. Similarly, for WS, a recent study by Rossi *et al.* found reduced monocarboxylate transporter 1 (MCT1) levels in the optic nerve of a WS mouse model, suggesting that the observed retinal neurodegeneration may in fact be caused by a lack of metabolic support from the oligodendrocytes. Furthermore, Samara *et al.* suggested that *WFS1* expression occurs predominantly in oligodendrocytes during early brain development [62]. This, together with neuroimaging studies showing abnormal white matter myelination in WS patients [40, 41, 61], led them to propose that WS could belong to a category of disorders characterized by ER stress-mediated myelination impairment [62]. Therefore, in this study, we aimed to better understand the role of oligodendroglia in WS pathogenesis and thereby lay the groundwork for the design of novel therapeutic strategies for neuroprotection in the WS CNS.

First, we found that *Wfs1* KO mice present with an age-dependent deterioration of visual function, starting from the age of 3 months and progressing to retinal ganglion cell dysfunction and impaired vision by the age of 7.5 months. Despite these overt functional deficits, we did not observe loss of retinal ganglion cells nor amacrine dopaminergic neurons in the retina of *Wfs1* KO mice up till the age of 7.5 months. This, combined with the unfolded protein response observed in the retina and optic nerve, suggests that we are looking at the earliest stages of disease, during which neuronal dysfunction is accumulating, but has not yet reached the tipping point towards cellular loss. Indeed, our findings confirm previous results in the same mouse model, in which neurodegeneration of the retinal ganglion cells was not seen at 8 months but manifested at 12 months [59], and mirror a study in WS patients, in which it was found that retinal ganglion cell axonal degeneration precedes cell body atrophy by about a decade [5]. Given the detrimental phenotype of this transgenic model, and applying humane end points, we limited our study to a maximum age of 7.5 months. At this age, we observed a mild and complex disease phenotype in the optic nerve, with alterations in axonal conduction strength combined with subtle thinning of the myelin sheath and a reduction of the OPC pool. Again, these findings are in line with a previous study in this *Wfs1*^ι1exon8^ mouse model [59], as well as in *Wfs1*^ι1exon2^ mouse and Wfs1^ι1exon5^ rat models [7, 53]. Novel findings in our study originate from *ex vivo* measurements of optic nerve CAPs, which allowed us to investigate a potential defect in the optic nerve while excluding the retinal ganglion cell bodies and synapses. These experiments uncovered a higher susceptibility to activity-induced conduction blocks in axons of *Wfs1* KO mice. These changes in axonal spiking capacity may arise from elevated K^+^ accumulation during firing or energy deficits in the axons. However, the speed of CAP peak recovery following high-frequency stimulation indicates that oligodendroglial potassium clearance [37] is not overtly affected in *Wfs1* KO animals.

Overall, the phenotype of the retina and optic nerve of the *Wfs1* KO mouse recapitulates one of the disease hallmarks of WS, i.e., optic neuropathy. Furthermore, the MRI study conducted in these mice confirms that the eye is a window to the brain and that manifestations of the disease in the brain stem, cerebellum and retinocollicular pathway recapitulate findings in WS patients (reviewed in [62]). Reduced volumes of the total brain, brainstem and cerebellum, as well as reduced fractional anisotropy values in the medulla, pons, cerebellum and superior colliculus, indicate that both grey and white matter are affected in the WS brain, and corroborate a myelin or oligodendrocyte defect.

Second, to further interrogate oligodendroglia function and assess whether any cell intrinsic defects may lead to primary oligodendroglia dysfunction and thereby increase vulnerability of the neurons, we performed a detailed and comprehensive assessment of cellular processes potentially affected in WS. Two pairs of *WFS1* mutant OPCs/pmOLs with concomitant isogenic controls were studied. A comparative transcriptomics study revealed that the transcriptome of the WS patient-derived OPCs/pmOLs was divergent, with few –mostly mitochondrial– differentially expressed genes in patient line 1, and patient line 2 having differentially expressed genes localising to the ER. This may reflect the phenotypic heterogeneity of the more than 200 mutations of the *WFS1* gene that have been described as causing the disease [50] and suggests that future studies should include several mutant cell lines/models. However, a comprehensive set of *in vitro* studies showed that these rather limited transcriptional changes do not translate into a MAM phenotype, as has been described for mutant *WFS1* iPSC-derived neurons [54, 75]. Specifically, the ER-mitochondria interactions, as measured with a proximity assay, were similar between mutant and isogenic oligodendroglia, and this was associated with normal ER Ca^2+^ storage, no increased ER stress susceptibility, normal mitochondrial activity and normal phosphatidylcholine, phosphatidylethanolamine, and phosphatidylinositol levels.

Other observations from the transcriptomics study include that both *WFS1* mutant cell lines had several top differentially expressed genes with a role in retinal homeostasis and associated with developmental disorders bearing striking similarities to WS, including syndromes with congenital deafness and retinal/optic nerve atrophy. Furthermore, two gene paralogs were found to be upregulated in patient line 2, namely *SERPINE1* and *SERPINE2*, and previous transcriptomics studies of pancreatic islets [20] and hippocampus and hypothalamus [21] of WFS1-deficient mice also revealed upregulated *SERPINA7* and *SERPINA10*. In the CNS, these serine protease inhibitors play roles that extend beyond their well-known function as inhibitors of fibrinolysis. SERPINE1 may play a role in neurodegeneration/neuroprotection via extracellular matrix remodelling, excitotoxicity, blood-brain barrier opening, and recruitment and activation of microglia [30], while secretion of SERPINE2 by astroglia/oligodendroglia at the axoglial junction is believed to regulate nodal length and myelin structure in the optic nerve and corpus callosum [13]. Aberrant expression of *SERPINE1* has been linked to neurodegenerative diseases, including Alzheimer’s [1, 27], Parkinson’s disease [57], and MS [28]. Serpinopathies, e.g., ALS (with SERPINE2 accumulation), are characterized by neuronal inclusion bodies that accumulate within the ER and elicit an ER overload response and cell death signalling, [10, 30, 43, 52]. *SERPINE1* and *SERPINE2* thus seem to be nodes connecting the pathways related to morphogenesis and development, wound healing and blood coagulation, regulation of cell death/apoptosis, and extracellular matrix, and even ER stress, that were observed to be differentially expressed in our transcriptomics analyses. Importantly, it has been proposed that targeting of SERPINs may constitute a clinically-relevant molecular approach to the therapy of neurodegenerative diseases associated with increased SERPIN levels [27]. Altogether, our transcriptomics analysis revealed differential expression of several disease-relevant targets. However, these transcriptional changes appeared not to result in a detrimental phenotype, as the cellular assays did not reveal any abnormalities in cellular functions known to be compromised in WS.

One striking yet also still puzzling finding from the transcriptomics study, is the large number of transcripts related to neuronal compartments, synapse, (synaptic) vesicles, and cell junction, and suppressed genes and gene networks that may be linked to axo-glial synaptic junctions. OPCs form synaptic complexes with neurons, and detect neuronal activity through various neurotransmitter receptors and ion channels [68]. At these axo-glial synapses, glutamate and GABA signaling play crucial roles in regulating OPC proliferation, oligodendrocyte maturation, and myelin formation [6, 25, 68, 77, 78]. Downregulated gene networks in the WS patient iPSC-derived OPCs/pmOLs, related to synapse maturation and ion channel clustering, suggest compromised axo-glial synaptic junctions. This bidirectional communication between neurons and oligodendroglia impacts OPC motility, proliferation, oligodendrocyte maturation, myelination, metabolic support for axons, and remyelination [46, 68]. One may speculate that the observed depletion of the OPC pool and abnormal myelin wrapping in 7.5-month-old *Wfs1* KO mice could be the result from abnormal neuron-OPC interactions.

The importance of neuron-glia communication points out one of the limitations of the current study, namely that we used monocultures of OPCs/pmOLs. Future studies with bi- or triculture systems of neurons, oligodendrocytes, and preferably also astrocytes, or organoids, are needed to study intercellular interactions in WS. These experiments are essential to clarify the cellular and molecular processes that, despite relatively normal cellular functions in the OPCs/pmOLs, lead to neuronal loss in WS. Another limitation may be that we used a population of oligodendrocyte lineage cells, of which only a small proportion expresses low levels of MBP and forms MBP^+^ extensions on electrospun fibers. These cells were defined as a mixed population of OPCs and pmOLs. Prolonged maturation up till 8-10 weeks, together with an axon or axon-like substrate for myelination, would be required for these cells to become mature, myelinating oligodendrocytes. As such, we cannot make any claims about a primary involvement of mature oligodendrocytes in WS, nor study processes related to myelination. Notably, the used OPCs/pmOLs are also likely to be less prone to ER stress as compared to myelinating oligodendrocytes. However, this disadvantage may have turned into a strength, and led to evidence for a role for OPCs rather than for fully mature oligodendrocytes at the axo-glial synapse. Intriguingly, further supporting the importance of OPCs for CNS function, neuronal loss was rescued and lactate production/release was restored in iPSC-derived motor neurons from ALS patients, by knockdown of the disease-causing gene *SOD1* in OPCs but not by knockdown in mature oligodendrocytes [14]. This underscores the importance of oligodendrocyte-lineage cells, especially OPCs, in neurodegeneration, and should be an incentive to further explore OPC proliferation, survival and differentiation dynamics in WS.

## 5. Conclusions

Following studies of the neurodegenerative phenotype of rodent WS models and WS patient iPSC-derived neurons, in this study, we focused on potential intrinsic defects in the oligodendroglia as a major contributor to WS pathogenesis. Our neuropathological and molecular findings suggest that cellular functions in *Wfs1* KO mouse oligodendrocytes and WS patient iPSC-derived OPCs/pmOLs are largely conserved. Given the cellular defects that have been previously observed in neurons in WS, and incentivised by the transcriptomic changes in the OPCs/pmOLs that may affect OPC-neuron synaptic junctions, and a depleted OPC pool in the optic nerve of *Wfs1* KO mice, we propose that future research should address neuron-glia communication to come to a better understanding of the pathogenesis of WS.

## 7. Declarations

### 7.1. Ethics approval and consent to participate

All animal experiments were performed according to the European directive 2010/63/EU and approved by the KU Leuven institutional ethics committee for animal research (project no. P068/2018). All human iPSC studies were done after approval by the KU Leuven institutional ethics committee (project no. S50354).

### 7.2. Consent for publication

Not applicable

### 7.3. Availability of data and material

The transcriptomics datasets generated and analysed during the current study are available in the GEO repository [GSE264222]. The lipidomics dataset is included as a supplement. All other datasets are available from the corresponding author on reasonable request.

### 7.6. Authors’ contributions

KA designed and performed experiments, analyzed data and contributed to the writing of the manuscript. MV, AN, EL, JVh, SB, JZ, JL, KN, TB, DDH, PB, ZL, TV, WG, AS, AN, EW, and UH designed and performed experiments, and analyzed data. MC analyzed data and contributed to the writing of the manuscript. MP, PA, LVDB, GB, AS, YCC, CV and LM designed the experiments, analyzed data, coordinated the study and edited the manuscript. LDG designed the experiments, analyzed data, coordinated the study and wrote the manuscript. All authors read and approved the final manuscript.

## Acknowledgements

We thank Prof. F. Urano (Washington University School of Medicine) for providing the Wolfram syndrome patient-derived iPSC lines; Lipometrix, Véronique Brouwers, Iene Kemps, Marijke Christiaens, Lut Noterdaeme, and Manuel Gutiérrez de Ravé Hidalgo for their excellent technical assistance; Lien Veys and Luca Masin for help with data analysis.

## 6. List of abbreviations

ALS: amyotrophic lateral sclerosis
BiP: immunoglobulin heavy chain binding protein
CAP: compound action potentials
CHOP: C/EBP homologous protein
CNS: central nervous system
DAPI: 4’,6-diamidino-2-phenylindole
EAAT2: excitatory amino acid transporter 2
ER: endoplasmic reticulum
GFAP: glial fibrillary acidic protein
IBA1: ionized calcium-binding adaptor molecule 1
iPSC: induced pluripotent stem cell
MAMs: mitochondria-associated endoplasmic reticulum membranes
MBP: myelin basic protein
MCT1: monocarboxylate transporter 1
MS: multiple sclerosis
OMM: oligodendrocyte maturation medium
OPCs: oligodendrocyte precursor cells
PDGFRα: platelet-derived growth factor receptor alpha
PFA: paraformaldehyde
PLA: proximity ligation assay
pmOLs: pre-myelinating oligodendrocytes
PTPIP51: protein tyrosine phosphatase interacting protein 51
RBPMS: RNA-binding protein with multiple splicing
ROI: region of interest
TBST: Tris-buffered saline with Tween-20
TH: thapsigargin
TM: tunicamycin
VAPB: vesicle-associated membrane protein
BWFS1: Wolfram syndrome 1
WS: Wolfram syndrome
XBP1: X-Box-Binding protein 1

