## Supplementary file 1 for "A deep phenotyping study in mouse and iPSC models to understand the role of oligodendroglia in optic neuropathy in Wolfram syndrome"

**Supplementary figures**

S
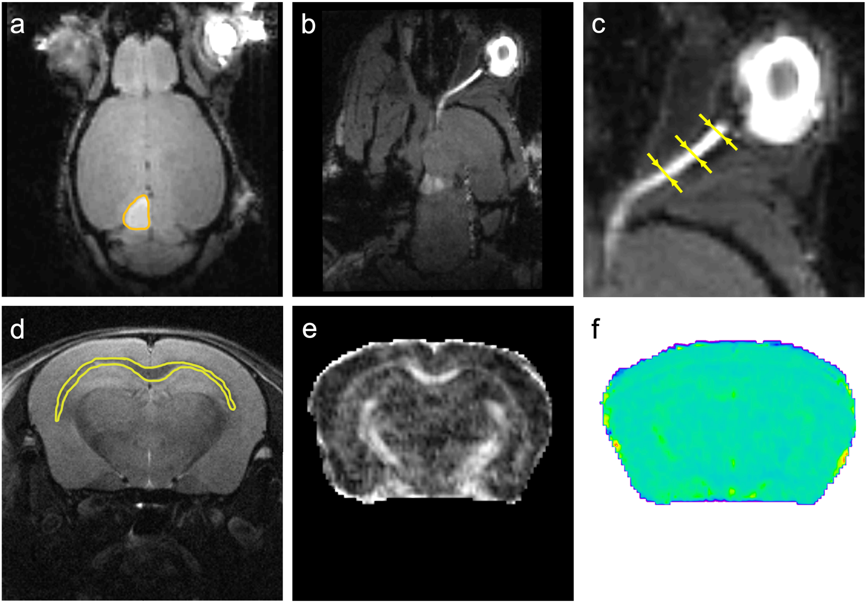

**Figure S1.** Illustration of parametric MR images and their analysis. **a-c** 3D T_1_-weighted MRI acquired 20 to 24 hours after intravitreal injection of MnCl_2_ into the right eye. Panel **a** shows hyperintense contrast in the right eye and superior colliculus in this horizontal image. The volume of the superior colliculus was calculated after manual delineation of the hyperintense region (orange line). Panel **b** shows the region of the optic nerve in the same 3D T_1_-weighted MRI as in (**a**). Panel **c** shows an enlarged section of (**b**). The yellow arrows indicate the locations where the diameter of the optical nerve was determined. The averaged diameter of the optic nerve at these three locations is shown in Figure 1h. **d** High-resolution anatomical MR image. Manual delineation of specific brain regions was used to determine their volumes as shown in Figure 4. The yellow line illustrates delineation of the corpus callosum (quantification shown in Figure 4d). **e-f** Delineations from 2D T_2_-weighted MR images were also projected on diffusion MR images for determining fractional anisotropy of diffusion (**e**) and mean diffusivity (**f**).

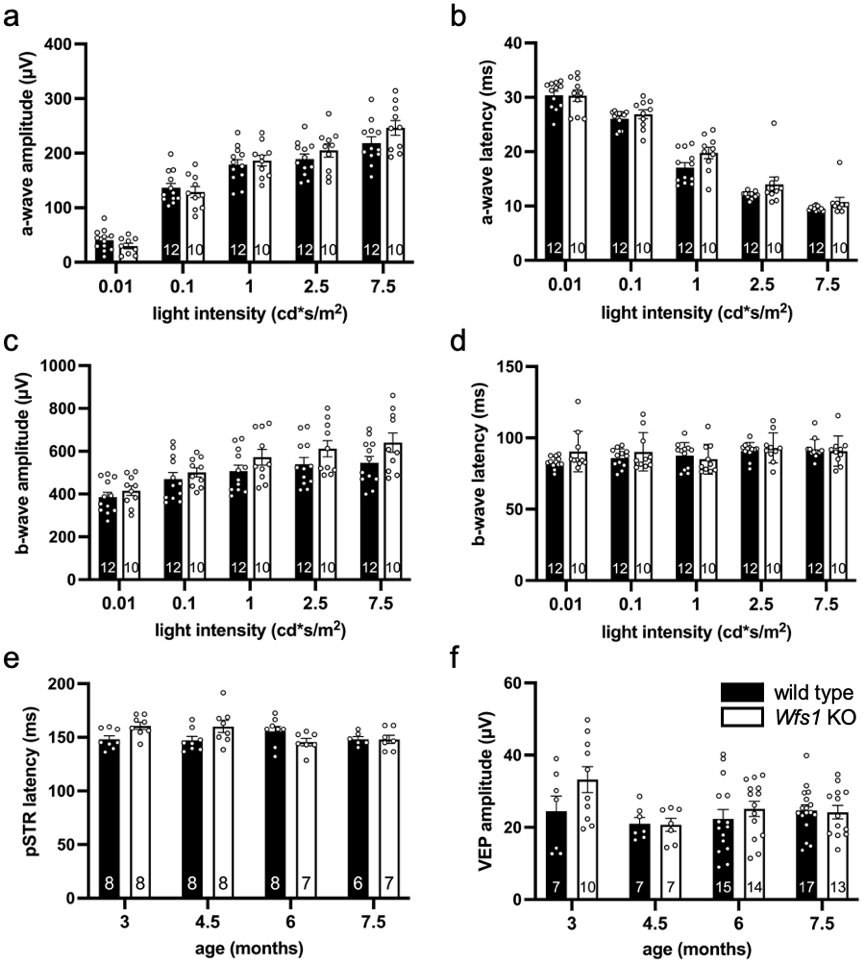

**Figure S2.** Electrophysiology data of *Wfs1* KO and wild type mice. **a-b** a-wave amplitude (**a**) and latency times (**b**) at 7.5 months of age. **c-d** b-wave wave amplitude (**c**) and latency times (**d**) at 7.5 months of age. **e** Positive scotopic threshold response latency at ages 3 till 7.5 months. **f** Visual evoked potentials amplitude at ages 3 till 7.5 months. No significant differences between genotypes were seen for any of the measurements, two-way ANOVA with Sidak’s multiple comparisons. Data presented as mean ± SEM, number of animals (N) depicted in the bar graphs.

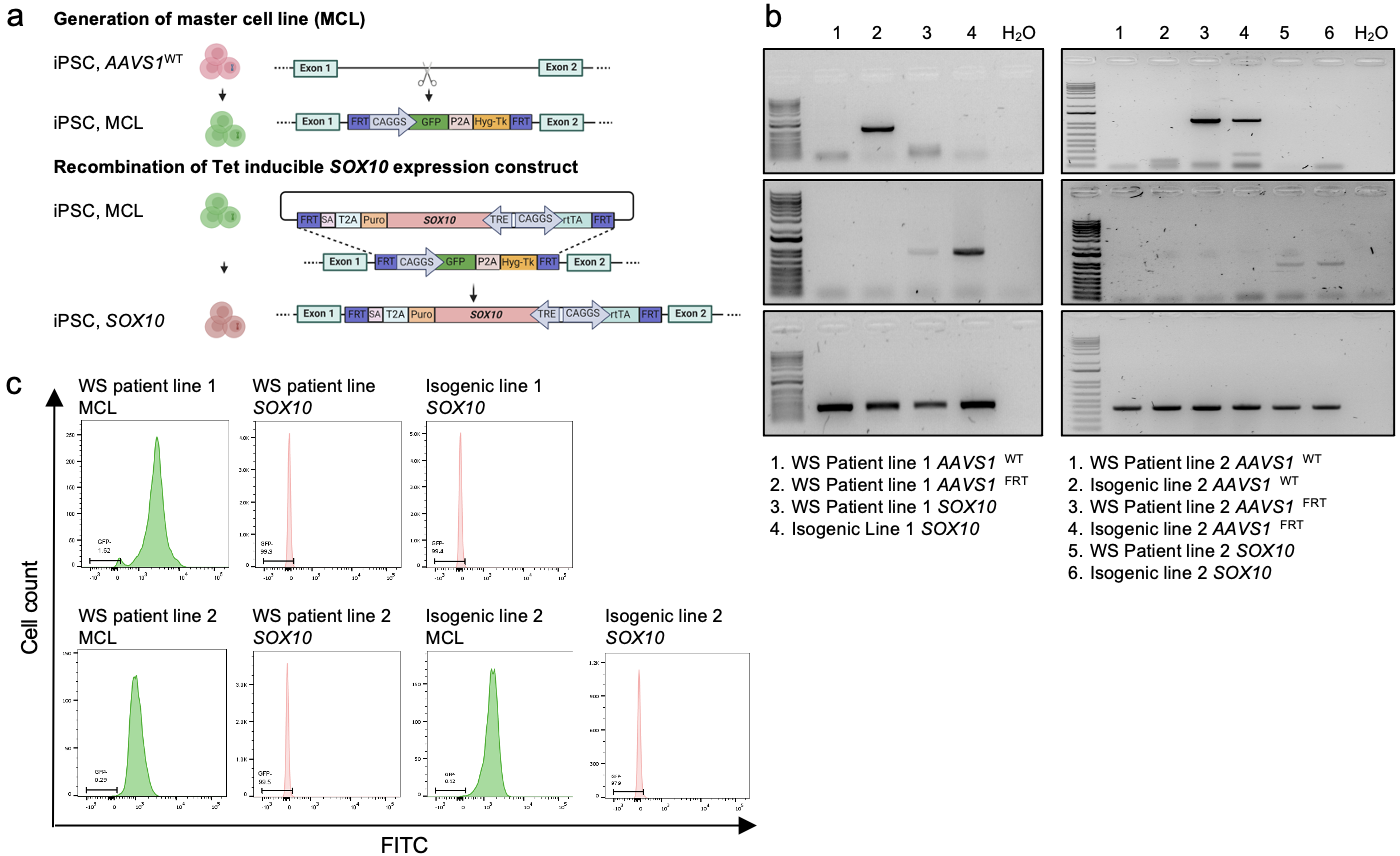

**Figure S3.** Integration of inducible *SOX10* in WS patient iPSC line 1 and 2, with their isogenic control iPSCs, via recombination mediated cassette exchange (RMCE). **a** Schematic representation of the RMCE process. First, a master cell line (MCL) was created by integration of a GFP-Hygro-TK cassette flanked by FRT-sites at the *AAVS1* locus by CRISPR nickase. Next, via RMCE, the GFP-Hygro-TK cassette was replaced with the construct encoding a TET-ON inducible *SOX10*. **b** Junction PCR analysis shows a clear distinct band representing presence of the correct cassette within the *AAVS1* locus, in MCLs (top gels) and SOX10 overexpression iPSC lines (middle gels) generated from WS patient lines and the isogenic pairs. MCLs contain the MCL constructs while RMCELs contain the RMCE constructs and no longer the MCL constructs. Non-modified iPSCs do not show any bands. Bottom gels represent wild type (WT) allele PCR, demonstrating that only one *AAVS1* allele is targeted in all the MCL and RMCELs. **c** Flow cytometric analysis of the MCL and *SOX10* overexpressing iPSC lines from WS patient lines 1 and 2 and isogenic controls. The first histogram for each cell line represents the MCL, revealing nearly 100% GFP positive cells. The second histogram represents *SOX10* overexpressing iPSC lines, which are nearly 100% GFP negative. For isogenic control 1, the *SOX10* iPSC line was directly generated from the WS patient *SOX10* iPSC line 1, therefore there is no MCL intermediate for this line.

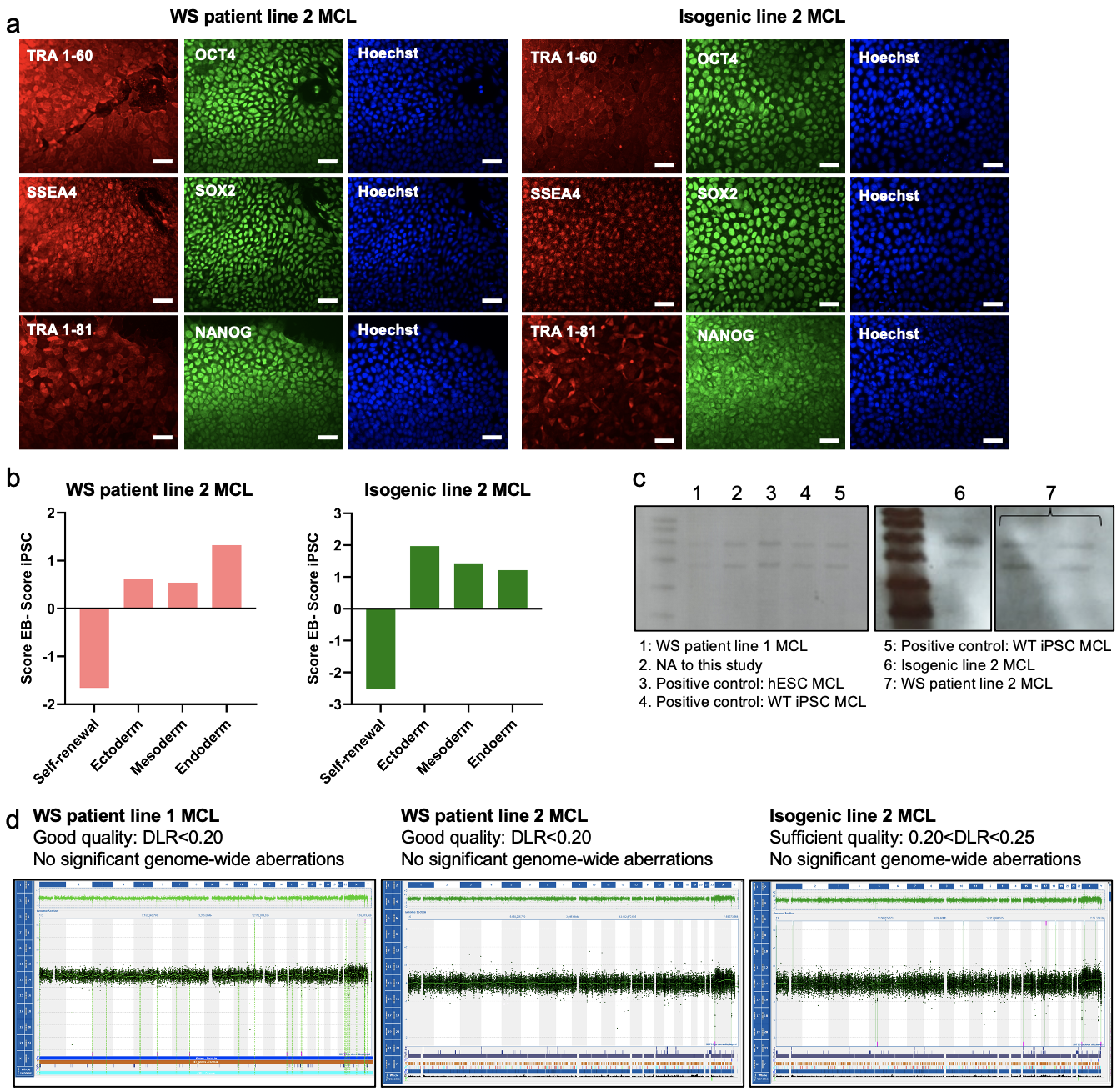

**Figure S4.** Quality control of master cell lines (MCL). **a** Immunocytochemical staining for pluripotency markers TRA-1-60, SSEA4, TRA-1-81, OCT4, SOX2 and NANOG in WS patient iPSC line 2 and the isogenic pair. **b** Transcripts for pluripotency, ectoderm, mesoderm and endodermal genes analyzed by ScoreCard®, of the embryoid bodies made from the MCLs of WS patient iPSC line 2 and its isogenic control. Scorecard analysis and pluripotency stainings were performed for WS patient line 1 in our previous study (Nami *et al*., 2021, Doi 10.1089/crispr.2021.0006). **c** Southern blot analysis of MCLs confirmed the absence of random integrations, and the correctly integrated FRT cassette in all iPSC lines. For isogenic control line 1, the *SOX10* iPSC line was directly generated from the WS patient *SOX10* iPSC line 1, therefore there is no data shown for this line. **d** Array-CGH results of the generated master cell lines show no chromosomal aberrations. Scale bar: 50 µm.

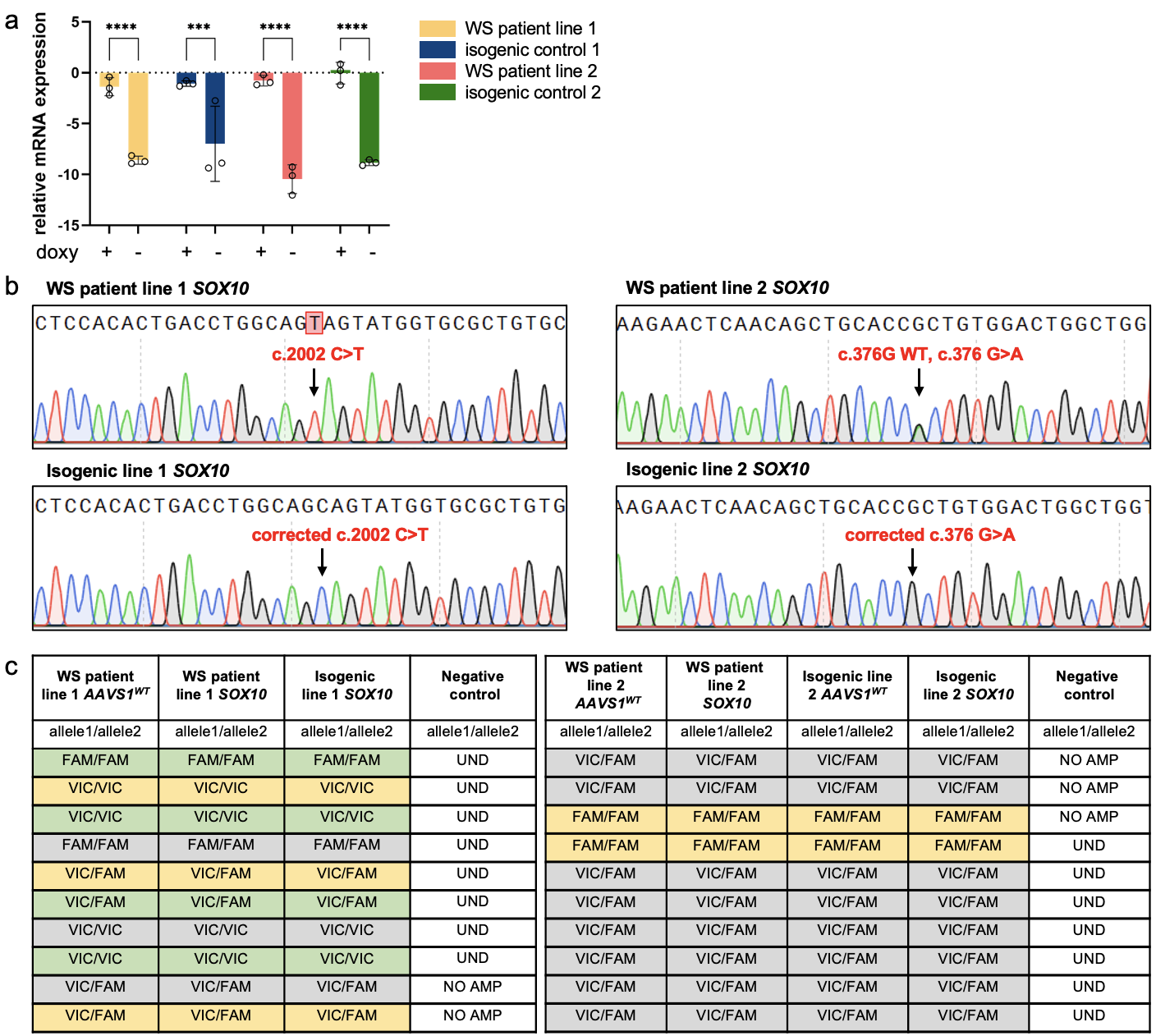

**Figure S5.** Characterization of inducible *SOX10* overexpressing WS patient and isogenic iPSC lines. **a** Expression of *SOX10* in WS patient and isogenic iPSC lines quantified by qRT-PCR. Unpaired t-test;  t_4_=12.85 for WS patient line 1, t_4_=2.776 for isogenic line 1, t_4_=11.16 for WS patient line 2, t_4_=13.82 for isogenic line 2, for without doxycycline *versus* with doxycycline. **b** Confirmation of *WFS1* mutations in *SOX10* WS patient lines and their correction in the isogenic controls via Sanger sequencing. Sequencing confirms the presence of the homozygous c.2002 C>T mutation in WS patient line 1, corrected in the isogenic control line 1; and heterozygous c.376 G>A mutation in WS patient line 2, corrected in isogenic control line 2. **c** SNP profiling (TaqMan® SNP Genotyping Assay) of WS patient and isogenic iPSC lines without any modification in the *AAVS1* locus, in comparison to inducible *SOX10* overexpressing iPSC lines and isogenic controls, revealed that the identity of the cell lines is identical.

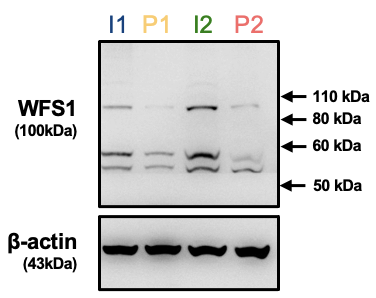

**Figure S6.** Western blot analysis of the Wolframin protein (WFS1) in WS patient and isogenic iPSC-derived OPCs/pmOLs confirms restoration of WFS1 expression in the isogenic OPCs/pmOLs of both lines. I1: isogenic line 1, P1: WS patient line 1, I2: isogenic line 2, P2: WS patient line 2.

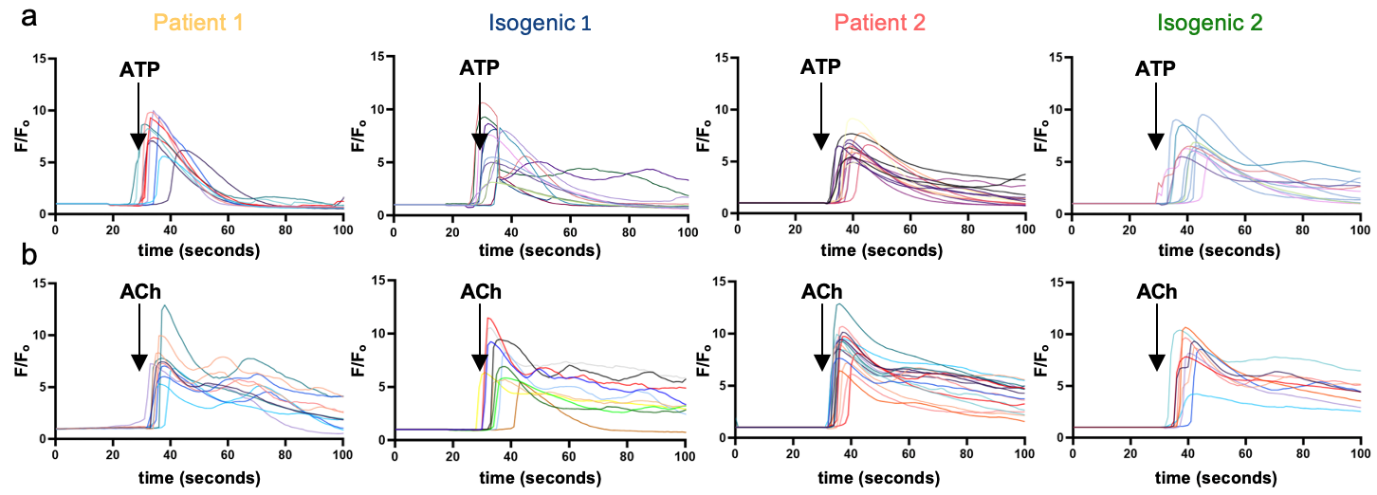

**Figure S7.** Representative Ca^2+^ traces following addition of 10 μM ATP (**a**) or 10 μM acetylcholine (ACh) (**b**), recorded from WS patient OPCs/pmOLs and their respective isogenic controls loaded with Cal-520. Time of ATP/ACh administration indicated with arrow. n=6 for WS patient line/isogenic control 1 and n=5 for WS patient line/isogenic control 2.

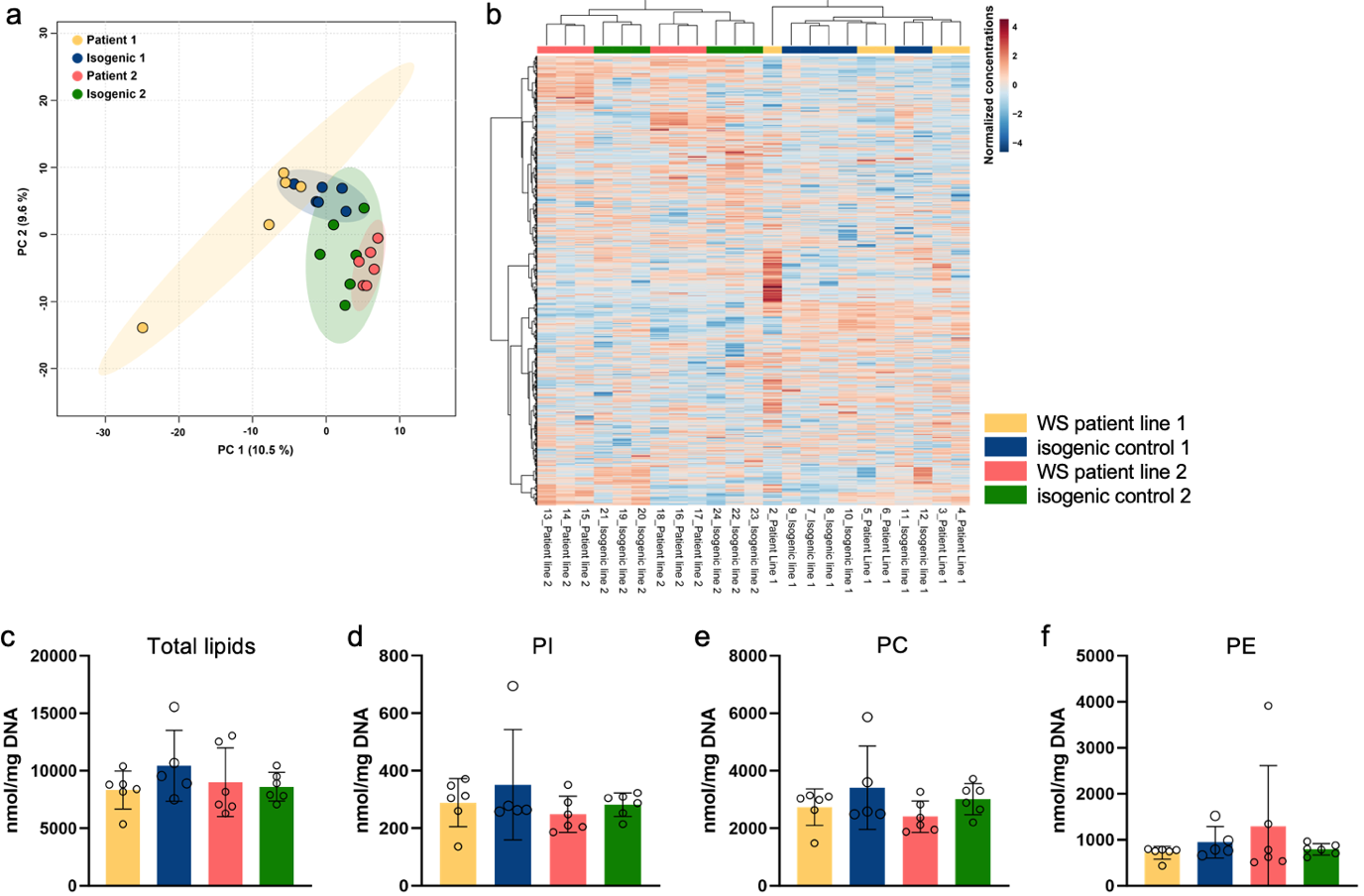

**Figure S8.** Mutations in the *WFS1* gene do not impact lipid metabolism in WS patient-derived OPCs/pmOLs compared to their isogenic controls. **a** PCA plot of lipidomics data in WS patient line 1 and 2 OPCs/pmOLs and their respective isogenic controls. **b** Heatmap showing similar lipidomics profiles in WS patient OPCs/pmOLs and isogenic controls. **c-f** Similar amounts of total lipids (**c**), phosphatidylinositol (PI) (**d**), phosphatidylcholine (PC) (**e**), and phosphatidylethanolamine (PE) (**f**) were found in WS patient and isogenic OPCs/pmOLs. Unpaired t-test; t_9_=1.461 for WS patient 1 *versus* isogenic 1 and t_10_=0.2973 for WS patient 2 *versus* isogenic 2 (**c**); t_9_=0.7250 for WS patient 1 *versus* isogenic 1 and t_10_=1.105 for WS patient 2 *versus* isogenic 2 (**d**); t_9_=1.038 for WS patient 1 *versus* isogenic 1 and t_10_=1.940 for WS patient 2 *versus* isogenic 2 (**e**); t_9_=1.487 for WS patient 1 *versus* isogenic 1 and t_10_=0.9080 for WS patient 2 *versus* isogenic 2 (**f**). n=6.

**Supplementary tables**

**Table S1.** Primer sequences

| *OLIG1* | GTACTCCTGCGTGTTAATGAGA  GCATCCAGTGTTCCCGATT |
| --- | --- |
| *OLIG2* | GACAAGCTAGGAGGCAGTGG  TCCGGCTCTGTCATTTGCTT |
| *SOX10* | CTTCATGGTGTGGGCTCAGG  CACTTTCGTTCAGCAGCCTC |
| *NKX2-2* | AAACCGTCCCAGCGTTAAT  CGGCTGACAATATCGCTACTC |
| *XBP1* | GGGTTAAGACAGCGCTTGGGGATGG  GGGAATCCATGGGGAGATGTTCTG |

**Table S2.** Statistics for data shown in Figure 7. Unpaired t-tests were used to compare the mRNA expression levels of several ER stress markers (*BIP, CHOP, IRE1A, ATF6* and *PERK*) in WS patient and isogenic OPCs/pmOLs, with/without tunicamycin (TM) or thapsigargin (TH) treatment (Figure 7a), and to compare parameters related to the bioenergetic profile of WS patient and isogenic OPCs/pmOLs (basal respiration, maximal respiration, ATP production, spare respiratory capacity and ECAR, measured by Seahorse assay) (Figure 7d).

| **Unpaired t-test** | **t-value (DF)** | **p-value** |
| --- | --- | --- |
| **BiP** | | |
| **WS patient line 1 vs. isogenic line 1** | | |
| Control | t_8_=0.5121 | p=0.6224 |
| TH | t_8_=0.9681 | p=0.3613 |
| TM | t_2_=0.4957 | p=0.6692 |
| **Isogenic 1** | | |
| Control vs. TH | t_9_=4.387 | p=0.0053 |
| Control vs. TM | t_9_=4.987 | p=0.0023 |
| TH vs. TM | t_9_=1.678 | p=0.3362 |
| **Patient 1** | | |
| Control vs. TH | t_9_=2.858 | p=0.0555 |
| Control vs. TM | t_9_=2.447 | p=0.1068 |
| TH vs. TM | t_9_=0.2862 | p=0.9895 |
| **WS patient line 2 vs. isogenic line 2** | | |
| Control | t_8_=1.897 | p=0.0944 |
| TH | t_8_=0.4393 | p=0.6721 |
| TM | t_2_=1.594 | p=0.2520 |
| **Isogenic 2** | | |
| Control vs. TH | t_9_=8.774 | p=0.0004 |
| Control vs. TM | t_9_=6.062 | p=0.0052 |
| TH vs. TM | t_9_=0.5709 | p=0.9149 |
| **Patient 2** | | |
| Control vs. TH | t_9_=1.177 | p=0.6935 |
| Control vs. TM | t_9_=3.259 | p=0.1063 |
| TH vs. TM | t_9_=2.370 | p=0.2660 |
| **CHOP** | | |
| **WS patient line 1 vs. isogenic line 1** | | |
| Control | t_8_=0.5063 | p=0.6263 |
| TH | t_8_=1.180 | p=0.2718 |
| TM | t_2_=0.6403 | p=0.5876 |
| **Isogenic 1** | | |
| Control vs. TH | t_9_=2.777 | p=0.0632 |
| Control vs. TM | t_9_=1.477 | p=0.4360 |
| TH vs. TM | t_9_=0.6220 | p=0.9085 |
| **Patient 1** | | |
| Control vs. TH | t_9_=6.946 | p=0.0002 |
| Control vs. TM | t_9_=4.624 | p=0.0037 |
| TH vs. TM | t_9_=0.6267 | p=0.9067 |
| **WS patient line 2 vs. isogenic line 2** | | |
| Control | t_8_=0.9529 | p=0.3686 |
| TH | t_8_=0.6137 | p=0.5565 |
| TM | t_2_=2.034 | p=0.1790 |
| **Isogenic 2** | | |
| Control vs. TH | t_9_=6.745 | p=0.0026 |
| Control vs. TM | t_9_=2.695 | p=0.1925 |
| TH vs. TM | t_9_=2.404 | p=0.2574 |
| **Patient 2** | | |
| Control vs. TH | t_18_=3.659 | p=0.0688 |
| Control vs. TM | t_18_=4.757 | p=0.0206 |
| TH vs. TM | t_18_=1.991 | p=0.3771 |
| **IRE1α** | | |
| **WS patient line 1 vs. isogenic line 1** | | |
| Control | t_8_=2.512 | p=0.0362 |
| TH | t_8_=0.8193 | p=0.4363 |
| TM | t_2_=1.373 | p=0.3034 |
| **Isogenic 1** | | |
| Control vs. TH | t_9_=5.241 | p=0.0016 |
| Control vs. TM | t_9_=2.606 | p=0.0829 |
| TH vs. TM | t_9_=1.355 | p=0.5038 |
| **Patient 1** | | |
| Control vs. TH | t_9_=7.005 | p=0.0002 |
| Control vs. TM | t_9_=5.662 | p=0.0009 |
| TH vs. TM | t_9_=0.3668 | p=0.9786 |
| **WS patient line 2 vs. isogenic line 2** | | |
| Control | t_8_=1.762 | p=0.1162 |
| TH | t_8_=1.317 | p=0.2244 |
| TM | t_2_=0.1957 | p=0.8629 |
| **Isogenic 2** | | |
| Control vs. TH | t_9_=7.440 | p=0.0014 |
| Control vs. TM | t_9_=5.609 | p=0.0083 |
| TH vs. TM | t_9_=0.01522 | P>0.9999 |
| **Patient 2** | | |
| Control vs. TH | t_9_=5.852 | p=0.0064 |
| Control vs. TM | t_9_=3.020 | p=0.1373 |
| TH vs. TM | t_9_=1.404 | p=0.5993 |
| **ATF6** | | |
| **WS patient line 1 vs. isogenic line 1** | | |
| Control | t_8_=0.9737 | p=0.3587 |
| TH | t_8_=0.2336 | p=0.8212 |
| TM | t_2_=1.200 | p=0.3529 |
| **Isogenic 1** | | |
| Control vs. TH | t_9_=2.317 | p=0.1310 |
| Control vs. TM | t_9_=0.4077 | p=0.9711 |
| TH vs. TM | t_9_=2.159 | p=0.1672 |
| **Patient 1** | | |
| Control vs. TH | t_9_=3.132 | p=0.0358 |
| Control vs. TM | t_9_=1.098 | p=0.6580 |
| TH vs. TM | t_9_=1.270 | p=0.5542 |
| **WS patient line 2 vs. isogenic line 2** | | |
| Control | t_8_=0.1422 | p=0.8904 |
| TH | t_8_=2.396 | p=0.0434 |
| TM | t_2_=2.346 | p=0.1436 |
| **Isogenic 2** | | |
| Control vs. TH | t_9_=1.771 | p=0.4547 |
| Control vs. TM | t_9_=0.8357 | p=0.8283 |
| TH vs. TM | t_9_=2.174 | p=0.3199 |
| **Patient 2** | | |
| Control vs. TH | t_9_=0.3298 | p=0.9706 |
| Control vs. TM | t_9_=1.670 | p=0.4929 |
| TH vs. TM | t_9_=1.420 | p=0.5927 |
| **PERK** | | |
| **WS patient line 1 vs. isogenic line 1** | | |
| Control | t_8_=0.8477 | p=0.4213 |
| TH | t_8_=0.07466 | p=0.9423 |
| TM | t_2_=2.727 | p=0.1123 |
| **Isogenic 1** | | |
| Control vs. TH | t_9_=7.850 | P<0.0001 |
| Control vs. TM | t_9_=3.149 | p=0.0349 |
| TH vs. TM | t_9_=2.786 | p=0.0623 |
| **Patient 1** | | |
| Control vs. TH | t_9_=3.507 | p=0.0198 |
| Control vs. TM | t_9_=2.453 | p=0.1058 |
| TH vs. TM | t_9_=0.1982 | p=0.9964 |
| **WS patient line 2 vs. isogenic line 2** | | |
| Control | t_8_=0.1886 | p=0.8551 |
| TH | t_8_=0.7318 | p=0.4852 |
| TM | t_2_=0.1480 | p=0.8959 |
| **Isogenic 2** | | |
| Control vs. TH | t_9_=6.737 | p=0.0026 |
| Control vs. TM | t_9_=1.771 | p=0.4545 |
| TH vs. TM | t_9_=3.322 | p=0.0994 |
| **Patient 2** | | |
| Control vs. TH | t_9_=3.782 | p=0.0601 |
| Control vs. TM | t_9_=1.279 | p=0.6513 |
| TH vs. TM | t_9_=1.580 | p=0.5279 |

| **Unpaired t-test** | | **t-value (DF)** | | **p-value** |
| --- | --- | --- | --- | --- |
| **Bioenergetic profile (Seahorse assay)** | | | | |
| **WS patient line 1 vs. isogenic line 1** | | | | |
| Basal respiration | t_8_=1.914 | | p=0.0919 | |
| Maximal respiration | t_8_=1.380 | | p=0.2050 | |
| ATP production | t_8_=2.193 | | p=0.0596 | |
| Spare respiratory capacity | t_8_=0.9893 | | p=0.3515 | |
| Extracellular acidification rate | t_8_=0.7187 | | p=0.4928 | |
| **WS patient line 2 vs. isogenic line 2** | | | | |
| Basal respiration | t_10_=0.6205 | | p=0.5488 | |
| Maximal respiration | t_10_=1.325 | | p=0.2146 | |
| ATP production | t_10_=0.2563 | | p=0.8029 | |
| Spare respiratory capacity | t_10_=2.121 | | p=0.0599 | |
| Extracellular acidification rate | t_4_=0.8944 | | p=0.4216 | |

**Table S3.** Statistics for data shown in Figure 8. Unpaired t-tests were used to compare parameters related to cytosolic Ca^2+^ release (normalized peak intensity and area under the curve, AUC) by WS patient and isogenic OPCs/pmOLs, stimulated by either ATP or acetylcholine (ACh).

| Unpaired t-test | t-score (DF) | Adjusted p-value |
| --- | --- | --- |
| **Cytosolic Ca^2+^ release** | | |
| **WS patient line 1 vs. isogenic line 1** | | |
| Peak-ATP | t_10_=0.3438 | p=0.7381 |
| Peak-ACh | t_10_=0.6146 | p=0.5526 |
| AUC-ATP | t=_10_2.278 | p=0.0459 |
| AUC-ACh | t_10_=0.2425 | p=0.8133 |
| **WS patient line 2 vs. isogenic line 2** | | |
| Peak-ATP | t_8_=1.359 | p=0.2113 |
| Peak-ACh | t_8_=0.4028 | p=0.6977 |
| AUC-ATP | t_8_=4.017 | p=0.0039 |
| AUC-ACh | t_8_=0.1075 | p=0.9170 |

**Supplementary information**

***Supplementary results***

*Generation of SOX10 overexpression iPSC lines*

Generation of *SOX10* overexpression iPSC lines was performed via the Recombinase-mediated Cassette Exchange (RMCE) approach [1]. This approach consists of two steps (Supplementary Figure S3a). In the first step, a master cell line is created via knock in of a selection cassette in the iPSC lines at the *AAVS1* locus. In the second step, this selection cassette is be exchanged with a *SOX10* overexpression cassette. In this study, the first step, namely the creation of the master cell lines, was performed using a selection cassette containing a hygromycin resistance gene, GFP and thymidine kinase, flanked by FRT sites (published in our previous study [2] and deposited in Addgene, plasmid #112666, pZ: F3-CAGGS GPHTK-F). Characterization of the master cell lines confirmed that the cells were GFP positive and contained the selection cassette, as detected by PCR (Supplementary Figure S3b and c). Pluripotency and the ability to differentiate to the three germ layer lineages of the master cell lines from patient 2 and the corresponding isogenic line were evaluated by staining and EB formation, followed by scorecards analysis. Results demonstrated that both of these two key characteristics of iPSC lines were maintained (Supplementary Figure S4a and b). Similar experiments for the patient 1 isogenic line, created by base editing, was published in our previous study [2]. Additionally, results of array-CGH showed no significant chromosomal aberrations in patient iPSC lines and isogenic genetically edited cells (Supplementary Figure S4d). In the second step, *SOX10* overexpression lines were created from master cell lines by exchanging this cassette with a *SOX10* overexpression cassette (pZ M2rtTA_CAGG TetON-Sox10**,** plasmid #115240) containing an inducible TET ON promoter expressing *SOX10* and a puromycin resistance gene. This second step was performed by nucleoporation of the master cell line with this construct, together with a Flippase-expressing vector. Successfully exchanged clones resistant to puromycin and fialuridine (FIAU; metabolized by TK to an active product that kills cells), which were also GFP negative by microscopy, were picked for further characterization. This included junction PCR, which confirmed the presence of the correct cassette at the *AAVS1* locus, and flow cytometry analysis, which confirmed the cells were GP negative, and qRT-PCR, which showed inducibility of *SOX10* cassette by doxycycline addition (Supplementary figure S3b and c, figure S5a). Additionally, Sanger sequencing results confirm presence of the mutations in the WS patient lines and the correction of the mutation in the isogenic lines (Supplementary Figure S5b). Furthermore, SNP profiling demonstrated the correct cell identity of the genome engineered lines (Supplementary Figure S5c).

***Supplementary methods***

*Stem cell culture, nucleoporation and colony isolation*

iPSCs were cultured on hESC qualified BD^TM^ matrigel (BD Biosciences) in E8 Flex medium (Thermo Fisher Scientific) in a humidified 5% CO_2_ incubator at 37°C, and passaged with 0.5mM EDTA (Gibco) in phosphate-buffered saline (PBS) when confluent. To perform nucleofection, cells were incubated at 37°C for 60’ in mTESR1 containing ROCKi (VWR Calbiochem), after which they were dissociated to single cell suspension by incubation with accutase (Sigma) for 10-15 minutes, based on the sizes of the colonies. Approximately 1x10^6^ cells were resuspended in 100μl nucleofection buffer and mixed with the DNA mixture, consisting of 2.5 μg of CRISPR/Cas9 nickase (Addgene plasmid #42335), 1 μg of the plasmid coding each guide RNA, and 5 μg of the donor plasmid, to create the master cell lines; or 2.5 μg Flippase, and 5 μg donor template to perform RMCE. Cells were transferred to a nucleofection cuvette and nucleofected using the Amaxa nucleofector II on setting F16. Afterwards, cells were immediately collected in mTESR1 supplemented with ROCKi, re-plated on BD^TM^ matrigel coated plates and placed in the incubator. The following day, medium was changed to E8 Flex supplemented with RevitaCell Supplement (100X, Gibco). Antibiotic selection was started when single cells grew to small colonies of 3-4 cells. To select for the master cell lines, cells were exposed to increasing concentrations of hygromycin (50-300 μg/ml), and to select for successful RMCE clones, cells were cultured with increasing concentrations of puromycin (100-300 ng/ml) and FIAU (1: 10000-1:5000; 0.5 mM). After ± 10-14 days, colonies were assessed by fluorescent microscopy to determine presence of GFP^+^ clones as an indication of successful master cell line generation, or GFP^-^ colonies as a result of successful RMCE generation. Selected colonies were picked manually using a colony picker tool (Fine Science Tools) under a microscope.

*Flow cytometry*

Flow cytometry was performed by dissociating the stem cell colonies to single cells as described before [2]. The cells were washed once and collected in PBS, and analyzed using the BD FACSCanto™ High Throughput Sampler. FCS files were analysed using flowjo v10.7.1 software.

*Junction PCR*

To perform junction PCR, genomic DNA from genome edited cells was extracted using the PureLink Genomic DNA kit (Invitrogen). Junction PCRs on *SOX10* overexpression lines were performed using the conditions outlined in Supplementary table 1.

*Southern blot*

To perform Southern blot, genomic DNA was extracted by using a DNeasy Blood & Tissue Kit (Qiagen). Genomic DNA was digested using NCOI enzyme (New England BioLabs) by incubating 7μl enzyme, 4μL Cutsmart buffer and 7μg genomic DNA in a total volume of 40 μL at 37˚C for 8 hours, and the inactivated at 85˚C for 20 minutes. A probe against the donor plasmid homology arms was generated by a PCR with Digoxigenin‐dNTPs (Roche) and Go Taq DNA polymerase (Promega). The PCR reaction conditions, and primer sequences are listed in Supplementary Table S4. Digested genomic DNA was loaded on a 0.7% agarose gel without Syber Safe and run long enough to ensure that the DNA was well separated. DNA was transferred from to a Hybond-N+ Nylon transfer membrane (Amersham) by capillary DNA transfer for 2 days. DNA was fixed by UV-crosslinking. Membrane hybridization and development were done using the DIG High Prime DNA Labelling and Detection Starter-Kit II (Roche) according to manufacturer’s instructions.

*Immunofluorescence staining*

Cells were fixed with 4% paraformaldehyde and incubated for 10 minutes at room temperature, and permeabilized with 0.1% Triton-X in PBS. Then, 30’ blocking was performed using 5% goat serum in PBS (Dako). The cells were stained overnight at 4°C with primary antibodies (Supplementary Table S5). After overnight incubation, cells were washed three times of 5 minutes with 0.1% Triton-X in PBS. Next, cells were incubated for 60 minutes at room temperature with the appropriate secondary antibodies, washed three times and then nuclei were stained with Hoechst 33342 (Sigma-Aldrich) 1:2000 diluted in PBS. Coverslips were mounted using Prolong Gold antifade reagent (Life Technologies) on superfrost microscopy slides (ThermoFisher Scientific). An AxioImager Z.1 fluorescence microscope (Zeiss) was used for taking images.

*Embryoid body (EB) formation and Scorecard analysis*

Approximately one million cells were detached using EDTA for 2 minutes and resuspended in 1 mL Essential 6 medium (Gibco) supplemented with 10µL RevitaCell, and plated into one well of Ultra-Low Attachment 24-well plates (Corning) for EB formation. EBs were maintained for 7 days with half medium changes every other day, after which they were collected for RNA isolation (GenElute^™^ Total RNA Purification Kit) followed by cDNA synthesis using the SuperScript™ III First-Strand Synthesis System kit (ThermoFisher scientific). Scorecard analysis was performed on cDNA of day 7 EBs, using TaqMan Gene Expression Master Mix in a TaqMan hPSC Scorecard® 384-well plate. Results were analysed by hPSC Scorecard® analysis software.

*Single nucleotide polymorphism (SNP) profiling*

SNP profiling was performed using TaqMan® SNP Genotyping Assay (Life Technologies). A custom made TaqMan® SNP Genotyping Assay (384-well plate), customized for genotyping SNPs, was loaded using 2x TaqMan^TM^ GTXpress^TM^ Master Mix 2.5 µl mixed with genomic DNA [4 ng/µl] 2.5 µl per well. Results were analysed by TaqMan Genotyper Software (Life Technologies).

*Array-CGH*

Genomic DNA samples were extracted using the PureLink Genomic DNA kit (Invitrogen). Array-CGH was performed at the Centrum Menselijke Erfelijkheid (CME) of the University Hospitals Leuven. Results were analysed using CytoSure interpret software.

**Supplementary Table S4.** Primers and PCR conditions for junction PCR and Southern blot probe production.

| **5’ junction PCR RMCEL** |  |
| --- | --- |
| Primers | CACTTTGAGCTCTACTGGCTTC  CATGTTAGAAGACTTCCTCTGC |
| Reagens | 2 μl goTaq grey buffer, 0.1 μl goTaq polymerase 1 μl MgCl_2_ 25 mM, 1 μl 2 mM dntp, 2μl primer mixes (2.5 μM), 50-100 ng genomic DNA in total volume of 10 μl |
| PCR conditions | 95°C, 5’ – [95°C, 30’’ – 72°C (‐0.5°C/cycle), 1’ 30’’] x 15 cycles – [95°C, 30’’ – 64°C, 30’’ – 72°C, 1’ 30’’] x 25 cycles – 72°C, 5’ |
| **AAVS1 WT allele PCR** |  |
| Primers | CTAGTCTTCTTCCTCCAACCCGG  GATCCTCTCTGGCTCCATCGTAAG |
| Reagens | 2 μl goTaq green buffer, 0.15 μl goTaq polymerase, 0.7 μl MgCl_2_ 25 mM, 1 μl 2 mM dntp, 1 μl from each primer (2.5 μM), 0.4 μl genomic DNA (50 ng/μl) in total volume of 10 μl |
| PCR conditions | 96°C, 5’ – [96°C, 30’’ ‐ 60°C, 30’’ ‐ 72°C, 1’] x 35 cycles – 72°C, 7’ |

**Supplementary Table S5.** List of primary and secondary antibodies used for pluripotency stainings.

| **Antibody** | **Company** | **Cat. No.** | **Dilution factor** |
| --- | --- | --- | --- |
| Rabbit Oct4 | Santa Cruz | Sc-9081 | 1:100 |
| Mouse TRA-1-60 (IgM) | Cell Signaling | 4746S | 1:200 |
| Rabbit Sox2 | Abcam | AB97959 | 1:500 |
| Mouse SSEA4 | Santa Cruz | sc-21704 | 1:200 |
| Rabbit Nanog | Thermo Scientific | PA1-097 | 1:300 |
| TRA-1-81(IgM) | Cell Signaling | 4745 | 1:200 |
| AF488 Goat anti-Rabbit | Life Technologies | A11034 | 1:500 |
| AF555 Goat anti-Mouse (IgM) | Life Technologies | A21426 | 1:500 |
| AF555 Goat anti-Mouse (IgG) | Life Technologies | A21424 | 1:500 |
